# Selective whole genome amplification as a tool to enrich specimens with low *Treponema pallidum* genomic DNA copies for whole genome sequencing

**DOI:** 10.1101/2021.07.09.451864

**Authors:** Charles M. Thurlow, Sandeep J. Joseph, Lilia Ganova-Raeva, Samantha S. Katz, Lara Pereira, Cheng Chen, Alyssa Debra, Kendra Vilfort, Kimberly Workowski, Stephanie E. Cohen, Hilary Reno, Yongcheng Sun, Mark Burroughs, Mili Sheth, Kai-Hua Chi, Damien Danavall, Susan S. Philip, Weiping Cao, Ellen N. Kersh, Allan Pillay

## Abstract

Downstream next generation sequencing (NGS) of the syphilis spirochete *Treponema pallidum* subspecies *pallidum* (*T. pallidum*) is hindered by low bacterial loads and the overwhelming presence of background metagenomic DNA in clinical specimens. In this study, we investigated selective whole genome amplification (SWGA) utilizing multiple displacement amplification (MDA) in conjunction with custom oligonucleotides with an increased specificity for the *T. pallidum* genome, and the capture and removal of CpG-methylated host DNA using the NEBNext^®^ Microbiome DNA Enrichment Kit followed by MDA with the REPLI-g Single Cell Kit as enrichment methods to improve the yields of *T. pallidum* DNA in isolates and lesion specimens from syphilis patients. Sequencing was performed using the Illumina MiSeq v2 500 cycle or NovaSeq 6000 SP platform. These two enrichment methods led to 93-98% genome coverage at 5 reads/site in 5 clinical specimens from the United States and rabbit propagated isolates, containing >14 *T. pallidum* genomic copies/μl of sample for SWGA and >129 genomic copies/μl for CpG methylation capture with MDA. Variant analysis using sequencing data derived from SWGA-enriched specimens, showed that all 5 clinical strains had the A2058G mutation associated with azithromycin resistance. SWGA is a robust method that allows direct whole genome sequencing (WGS) of specimens containing very low numbers of *T. pallidum*, which have been challenging until now.

**Importance:** Syphilis is a sexually transmitted, disseminated acute and chronic infection caused by the bacterial pathogen *Treponema pallidum* subspecies *pallidum*. Primary syphilis typically presents as single or multiple mucocutaneous lesions, and if left untreated, can progress through multiple stages with varied clinical manifestations. Molecular studies often rely on direct amplification of DNA sequences from clinical specimens; however, this can be impacted by inadequate samples due to disease progression or timing of patients seeking clinical care. While genotyping has provided important data on circulating strains over the past two decades, WGS data is needed to better understand strain diversity, perform evolutionary tracing, and monitor antimicrobial resistance markers. The significance of our research is the development of a SWGA DNA enrichment method that expands the range of clinical specimens that can be directly sequenced to include samples with low numbers of *T. pallidum*.

## Introduction

Syphilis, caused by *Treponema pallidum* subspecies *pallidum* (hereafter referred to as *T. pallidum*), is steadily increasing in the United States (U.S.). In 2019, 38,992 cases (11.9 per 100,000 people) of primary and secondary (P&S) syphilis and 1,870 cases (48.5 per 100,000 live births) of congenital syphilis were reported to the CDC; representing a 167.2% increase in P&S syphilis rates since 2010 and a 291.1% increase in congenital syphilis since 2015 (1). Penicillin has been the drug of choice for treating all stages of syphilis; however, azithromycin has been used as an alternative to penicillin for treating early syphilis or contacts of syphilis cases worldwide. Macrolide-resistant *T. pallidum* strains associated with two mutations (A2508G, A2509G) in the 23S rRNA genes, have since been reported worldwide (2–5). While macrolides are not recommended as first line treatment in many countries, periodic monitoring is useful to determine the prevalence of resistant strains (6).

Molecular studies on contemporary *T. pallidum* strains in the U.S. remains challenging due to the limited number of sequenced whole genomes directly from clinical specimens. Strain diversity has been gleaned from molecular epidemiology studies, which are based on a few genetic loci, but may not be representative of the entire *T. pallidum* genome (7–10). In addition, studies have relied primarily on strains propagated in rabbits or DNA amplified directly from clinical specimens, because *T. pallidum* cannot be grown on routine laboratory media. However, advances have been made with *in vitro* tissue culture and the propagation of *T. pallidum* in rabbits from cryopreserved genital lesion specimens, which may make routine culture directly from clinical specimens a possibility in the near future (11–13).

Metagenomic shotgun sequencing approaches have made significant advances in recent years with sequence data being used for pathogen detection, whole genome-based typing, and antimicrobial resistance marker detection, in addition to phylogenetic analyses (14–16). However, direct whole genome sequencing (WGS) of *T. pallidum* from clinical specimens and rabbit isolates can be problematic due to bacterial genomic DNA (gDNA) being outweighed by either human or rabbit DNA, respectively. Several DNA enrichment methods have been described for *T. pallidum* including RNA bait capture techniques, methyl-directed enrichment using the restriction nuclease DpnI, and pooled whole genome amplification (17–22). These methods have generated *T. pallidum-specific* WGS data from over 700 metagenomic samples; however, sequencing specimens with low numbers of *T. pallidum* remains challenging. Therefore, new approaches that would enable sequencing of samples with low bacterial loads are needed. Host DNA removal by 5’-C-phosphate-G-3’ (CpG) methylated capture and selective whole genome amplification (SWGA), which allows selective amplification of gDNA of the species of interest compared to host DNA, have been successfully used for enriching bacterial gDNA in metagenomic samples (23–27); however, these methods have not been applied to *T. pallidum*.

In this study, we describe a DNA enrichment method based on selective whole genome amplification (SWGA) using multiple displacement amplification (MDA) and custom primers that enables WGS of clinical specimens with very low genomic copies of *T. pallidum*. We also investigated an alternative method, the NEBNext^®^ Microbiome DNA Enrichment Kit, that uses CpG methylated capture of host DNA followed by MDA with the REPLI-g Single Cell Kit.

## Results

### Real-time qPCR on clinical specimens and spiked samples

A total of 15 clinical specimens were included in this study (Table 1). DNAs were extracted using standard or large-scale extraction protocols and, *T. pallidum* gDNA (copies/μl of extract) was estimated using qPCR targeting the *polA* gene. As shown in Table 1, 3 specimens had >100 *T. pallidum* gDNA copies/μl and the remaining 12 specimens had < 32 copies/μl. PCR amplification of the human RNase P gene (RNP) was used to estimate the concentration of human gDNA in each sample (28). RNP Ct values in the specimens ranged from 22.61 (highest concentration of RNP) to 38.32 (lowest concentration of RNP) (Table 1).

**Table 1.**
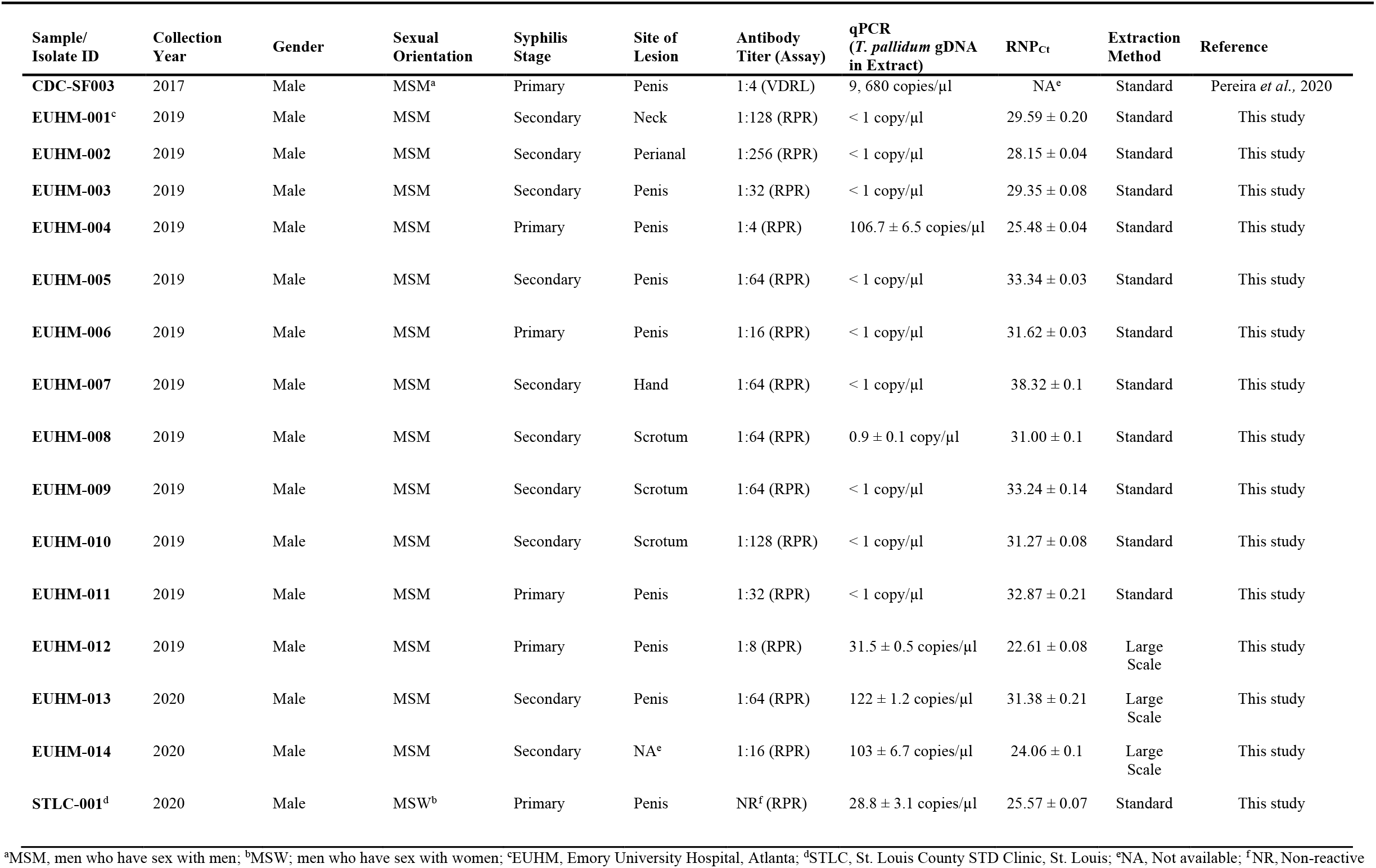
Clinical and laboratory data for specimens and the *T. pallidum* isolate.

To determine the limit of detection (LoD) for each enrichment method, spiked samples were composed of a 10-fold dilution series of *T. pallidum* Nichols DNA against a background of a constant concentration of human DNA. An RNP_Ct_ of 25.48, corresponding to the lowest C_t_ value among specimens extracted with the standard protocol, was targeted as the cut-off for all the spiked samples. Based on qPCR, the copy number of the undiluted spiked sample was estimated at 11,066.17 ± 364.60 gDNA copies/μl with an average RNP_Ct_ value of 25.07 ± 0.04 (Table S1). The dilution series generated from this spiked sample averaged 1,016 ± 9.41 down to 1.57 ± 0.12 *T. pallidum* gDNA copies/μl. The RNP_Ct_ values averaged 25.02 ± 0.03 amongst all samples, with no significant difference observed amongst the RNP_Ct_ values for each dilution in the series (P > 0.3; Table S1).

### NEBNext microbiome enrichment with MDA

To determine the minimum *T. pallidum* copy number input required to generate adequate sequencing coverage, each of the replicate 10-fold diluted spiked samples were enriched with the NEB Microbiome Enrichment Kit with subsequent REPLI-g Single Cell MDA (hereafter referred to as NEB+MDA). We observed an increase in *T. pallidum* gDNA, determined by qPCR, in all spiked samples after enrichment (Fig. 1, Table S1). The enriched spiked samples indicated an average of 6.67 × 10^6^ ± 2.74 × 10^5^ to 964 ± 574.23 gDNA copies/μl in the undiluted to 1:10,000 diluted samples, resulting in a 482 to 995.09X enrichment (Table S1). Upon comparing the average RNP_Ct_ of each dilution in the series, the enriched samples indicated 29.28 ± 1.07 to 31.42 ± 0.45 for the neat to 1:10,000 dilution (Table S1). The RNP_Ct_ value of each enriched sample in the dilution series were not significantly different from one another, with an average RNP_Ct_ = 30.64 ± 0.33 for all dilutions in the series (P = 0.22).

**Fig 1.**
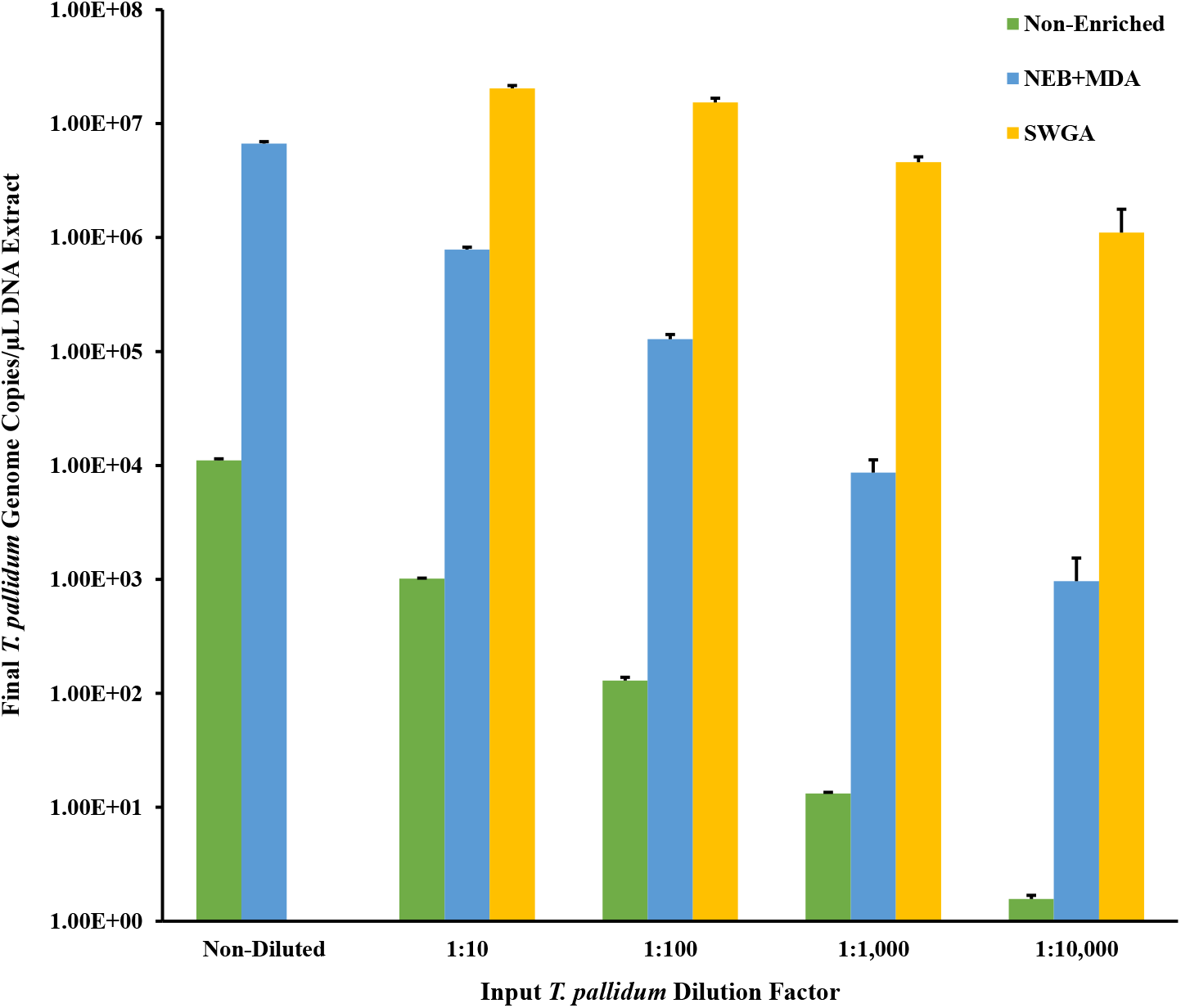
*T. pallidum* gDNA copies/μl for the 10-fold dilution series spiked samples enriched by the NEB+MDA or SWGA. Spiked samples were composed of a 10-fold dilution series of *T. pallidum* Nichols DNA and a constant concentration of human DNA. *T. pallidum* genome DNA (copies/μl of DNA extract) in samples pre- and post-enrichment was estimated using PCR targeting the *polA* gene and are shown in the bar graph. The y-axis has been log10 scaled for depiction of the Non-Enriched dilution series. Error bars represent standard error among three replicate enriched *T. pallidum* samples.

After enriching with NEB+MDA, the average percent of *T. pallidum* DNA in the total DNA ranged from 2.33% ± 0.10 to 0.0004% ± 0.0003% for the neat to 1: 10,000 dilutions samples, resulting in up to a 26.12-fold increase in the percent of *T. pallidum* DNA amongst all enriched replicates (Fig. 2). With the exception of enriched samples from the 1:100 and 1:1,000 dilutions, we observed that increasing the *T. pallidum* input gDNA copy number 10-fold resulted in a significant increase in the total DNA belonging to *T. pallidum* post-enrichment (P = 0.06 and P < 0.05, respectively).

**Fig 2.**
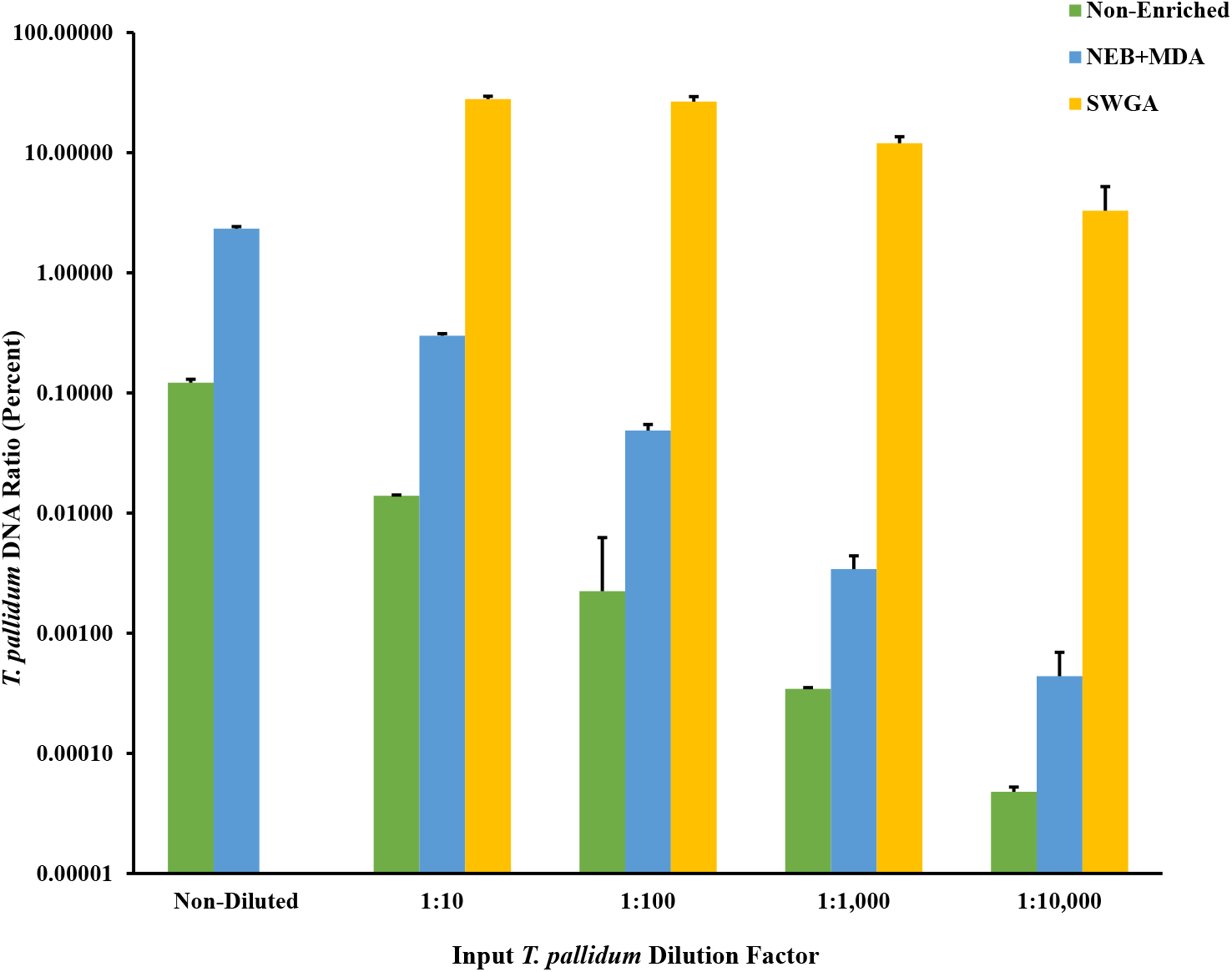
Relative percent *T. pallidum* Nichols DNA in total DNA for Non-Enriched, NEB+MDA, and SWGA enriched spiked samples. Spiked samples were composed of a 10-fold dilution series of *T. pallidum* Nichols DNA and a constant concentration of human DNA. Percent *T. pallidum* DNA in total DNA was calculated based on the input DNA concentration and gDNA copies/μl (Non-Enriched), and the DNA concentration and gDNA copies/μl for the Nichols-spiked samples post-enrichment (NEB+MDA or SWGA). Genome copies were estimated from measured *T. pallidum polA* copies/μl of DNA extract. The y-axis is log10 scaled for depiction of the Non-Enriched dilution series. Error bars represent standard error among three replicate samples.

To confirm if the increase in *T. pallidum* gDNA copies correlated with the increase in genome coverage, sequencing data derived from samples enriched by NEB+MDA were mapped against the *T. pallidum* Nichols reference genome (*NC_021490.2*). The genome sequencing data showed 0.01% to 10.52% of the quality-controlled reads binned as *T. pallidum*, along with a mean read mapping depth to *T. pallidum* Nichols reference genome ranging from 0.51 to 501.75X. An average percent coverage exceeding 97.29% across the Nichols reference genome with at least 5 reads mapped per site (5X) for the neat to 1:100 diluted samples, was observed among the NEB+MDA enriched samples (Fig.3A; Table S1 and Fig. S1). At the same time, for a higher coverage of at least 10 reads mapped per nucleotide (10X), the 1:100 diluted samples had an average percentage coverage of 84.14% while neat and 1:10 dilution samples covered at 99.99% across the reference genome. A sharp decline in coverage was observed in the enriched 1:1,000 and 1,10,000 diluted samples. With the QC criteria for efficiency set at ≥90% at ?5X read depth, samples sequenced post NEB+MDA enrichment had a LoD of 129 *T. pallidum* gDNA copies/μl (Fig. 3A; Table S1 and Fig. S1). A comparison of all the genome data generated from the serially diluted samples and the non-enriched *T. pallidum* Nichols control using the NEB+MDA protocol revealed no genetic variants, verifying that sequencing errors were not introduced during whole genome amplification. NEB+MDA was subsequently used to enrich CDC-SF003, a recently propagated clinical isolate (11), with 2.39 × 10^6^ ± 1.35 × 10^5^ gDNA copies/μl of DNA extract observed post-enrichment. We observed that 1.06% of the total DNA belonged to *T. pallidum* post enrichment and 3.29% of the host removed quality-controlled sequencing reads were classified as *T. pallidum*. Sequencing indicated a 98.60% coverage across the *T. pallidum* SS14 reference genome (*NC_021508.1*) at 5X read depth with a mean mapping depth of 46.43 (Fig. 4; Table 2 and Fig. S2).

**Fig 3.**
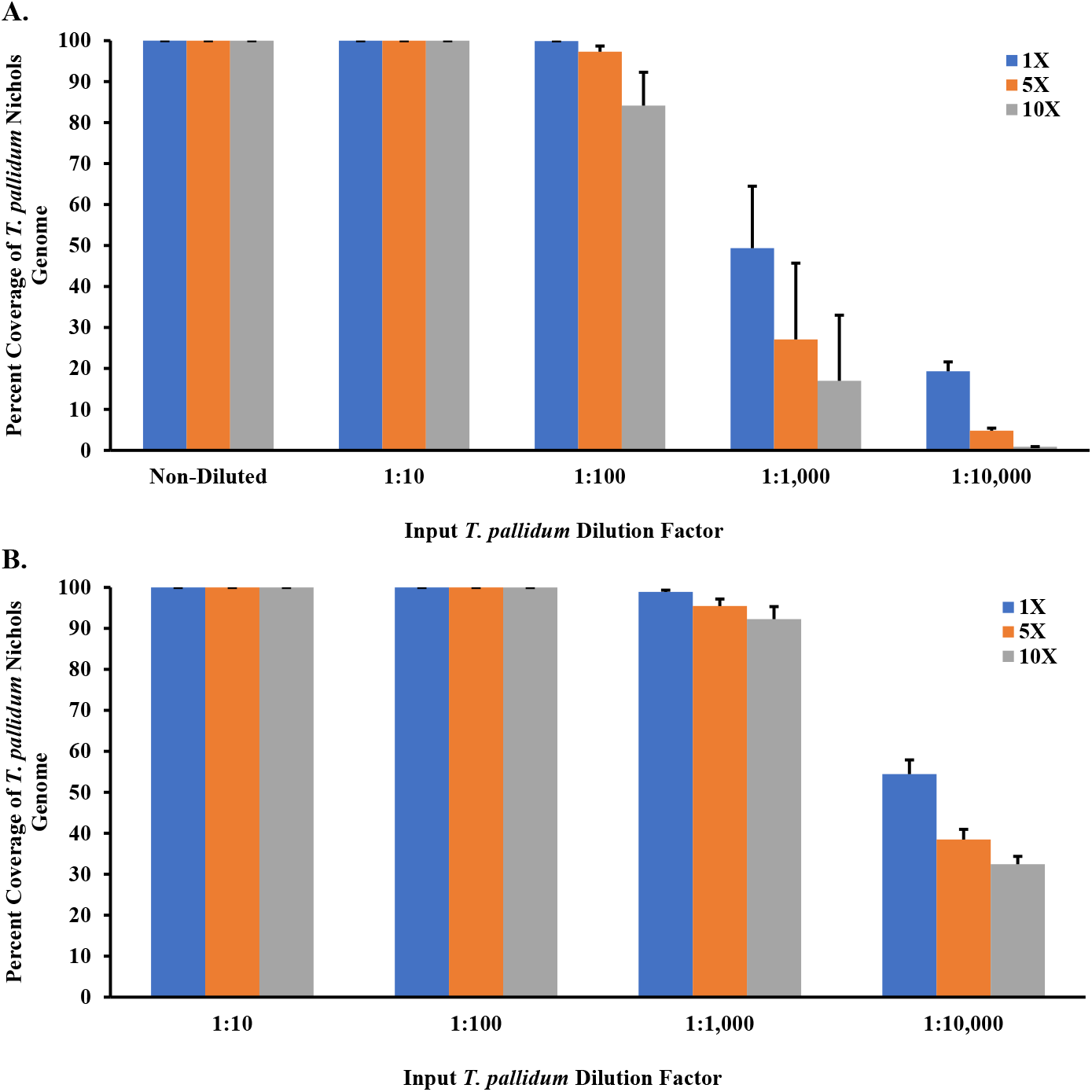
Percent coverage of sequencing reads of enriched *T. pallidum* Nichols spiked samples. Treponemal reads with at least 1 read mapped per site (1X) against the *T. pallidum* subsp. *pallidum* Nichols reference genome (*NC_000919.1*) and percent coverage of *T. pallidum* genome is shown. (A) Sequencing reads of samples enriched using the NEB+MDA method. (B) Sequencing reads of samples enriched using SWGA. All samples were sequenced using the Illumina NovaSeq 6000 platform. Error bars represent standard error between the mapped reads derived from three replicate enriched Nichols samples.

**Fig 4.**
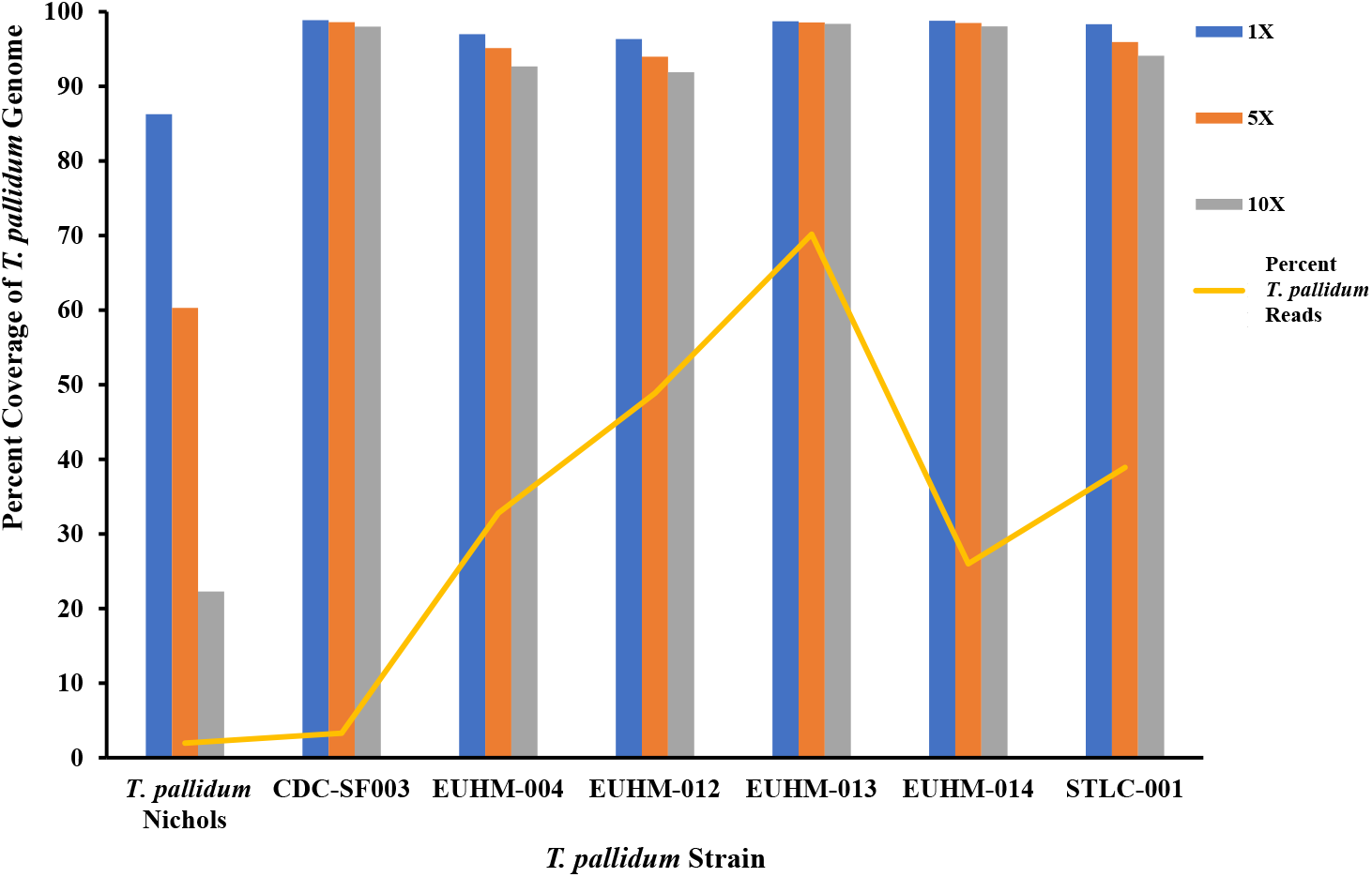
Percent coverage of *T. pallidum* isolates and clinical specimens. All samples were sequenced using the Illumina MiSeq v2 (500 cycle) platform. Percent of *T. pallidum* reads are derived from down selected *T. pallidum* reads. Prefiltered reads for Nichols-CDC were mapped to the Nichols reference genome (*NC_000919.1*). The prefiltered reads in all clinical isolates and specimens were mapped against the SS14 reference genome (*NC_021508.1*).

**Table 2.**
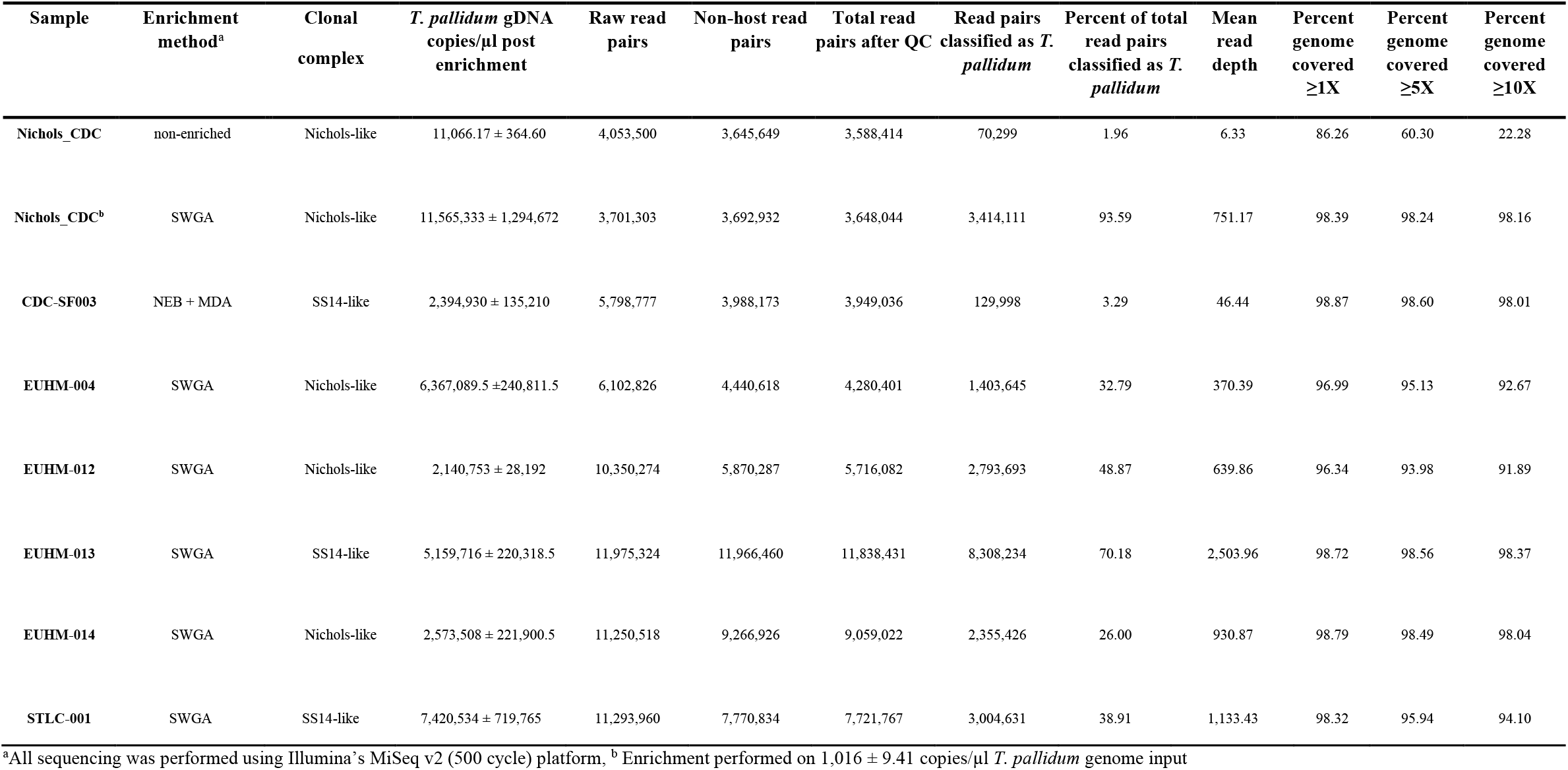
Sequencing percent coverage for the Nichols isolates, clinical isolate CDC-SF003, and clinical specimens across the *T. pallidum* reference genome.

### SWGA enrichment of *T. pallidum* Nichols

While NEB+MDA was useful for enriching samples with >129 *T. pallidum* gDNA copies/μl, we sought an alternative method for enriching clinical specimens with lower *T. pallidum* gDNA copies (Table 1). Selective whole genome amplification (hereafter referred to as SWGA) was chosen based on its success in other bacteria (25–27). To determine an optimal primer set for enriching *T. pallidum* during SWGA, a total of 12 primer sets were tested using Equiphi29 MDA (Thermo Fisher Scientific, Waltham, MA; Table S2-S3) and the 1:100 diluted Nichols DNA sample (~129 gDNA copies/μl) (Table 1; Table S4). The efficacy of the SWGA primer sets varied, resulting in DNA enrichment differences between 6.86 to 1.16 × 10^5^ times (Fig. 5A) with 7 of the 12 primer sets (SWGA Pal 2, 4, 5, 9, 10, 11, and 12) and a >10,000-fold increase in gDNA copy number. SWGA Pal 9 and Pal 11 gave the highest enrichment at 1.13 × 10^5^, and 1.16 × 10^5^ times, respectively (Table S4). The difference observed between Pal 9 and Pal 11 in the *T. pallidum* gDNA copy number and relative percent DNA belonging to *T. pallidum* was not significantly different and Pal 11 was selected for testing the SWGA limit of detection (P > 0.1; Fig. 5 and Table S4).

**Fig 5.**
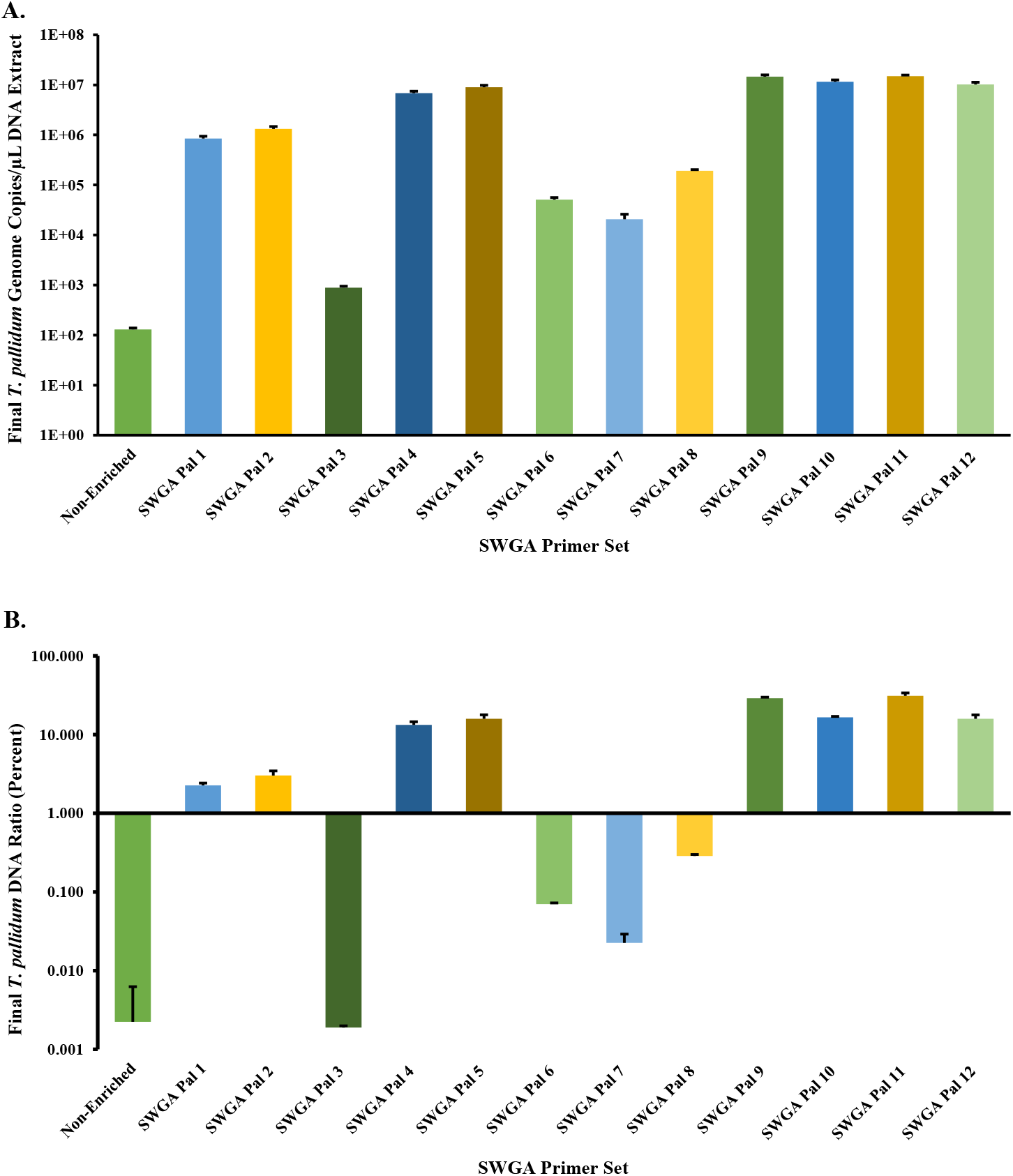
SWGA primer set validation. (A) *T. pallidum* gDNA copies/μl for the Nichols spiked sample (1:100 diluted) enriched with each SWGA primer set (1 to 12). (B) Relative percent *T. pallidum* DNA for the Nichols spiked sample (1:100 dilution) enriched with each SWGA primer set. Percent *T. pallidum* DNA was calculated based on the input DNA concentration and gDNA copies/μl for the Nichols mock samples post-SWGA enrichment. The spiked sample contained purified human gDNA, and the *T. pallidum* genome copies were derived from qPCR measured *T. pallidum polA* copies/μl of DNA extract. The input *T. pallidum* gDNA copies/μl of DNA is displayed as Non-Enriched. The y-axis is log10 scaled in each panel for depiction of the relative percent *T. pallidum* post-enrichment with each primer set. Error bars represent standard error among three replicate Nichols samples.

To determine the SWGA Pal 11 primer set’s LoD and enrichment for *T. pallidum*, SWGA was performed in triplicate on the 10-fold dilution series, except the neat sample. We observed a marked increase in *T. pallidum* gDNA copy number in every dilution in the series post enrichment (Fig. 1; Table S1). The gDNA copy number ranged from 1.11 × 10^6^ *±* 6.68 × 10^5^ for the 1:10,000 dilution to 2.04 × 10^7^ *±* 1.20 × 10^7^ in the 1:10 dilution (Table S1). When compared to the input copy number, this was a 2.01 × 10^4^-fold to 5.53 × 10^5^-fold increase in the enriched samples, from 1:10 −1:10,000 dilution, respectively. Upon comparing the average RNP_Ct_ of each dilution in the series, the SWGA enriched samples indicated a 29.36 ± 0.37 to 28.65 ± 0.16 for the 1:10 −1:10,000 dilution, respectively (Table S1). The average RNP_Ct_ at each 10-fold increase in *T. pallidum* concentration was not significantly different from one another (P > 0.1); however, by increasing the *T. pallidum* gDNA input 100-fold, we observed a significant decrease in RNP concentration (P < 0.03).

After enriching with SWGA, we observed that dilutions ranging from 1:10 to 1:10,000 held 27.93% ± 1.57% to 3.29% ± 1.93% of the total DNA belonging to *T. pallidum*, respectively (Fig. 2). This reflected up to a 1.63 × 10^5^-fold increase in the relative *T. pallidum* amongst all replicate SWGA enriched samples compared to unenriched samples. All samples were significantly increased in their relative *T. pallidum* DNA when compared to their respective inputs (P < 0.0001). While there was observed deviations in the percent DNA between replicates, the 1:10,000 diluted replicates still yielded a 28.40-fold ± 17.71-fold increase in DNA belonging to *T. pallidum* post-SWGA when compared to the non-enriched neat dilution (P < 0.0001).

Genome sequencing data derived from the SWGA enriched Nichols samples showed 0.98% to 78.05% of the quality-controlled reads binned as *T. pallidum*, along with a mean mapping depth to *T. pallidum* Nichols reference genome ranging from 65.82 to 4.89 × 10^3^. An average percent coverage of 98.67% ± 0.005% for the 1:10 diluted samples down to 96.15% ± 0.082% in the 1:1,000 diluted samples across the Nichols genome at 5X read depth was observed among the SWGA enriched samples. A sharp decline in coverage was observed at the 1:10,000 dilution (Fig. 3B; Table S1 and Fig. S3). Comparing NEB+MDA and SWGA for enrichment, we observed that SWGA consistently produced higher relative *T. pallidum* DNA in all 10-fold diluted samples (P < 0.01). In addition, there was a sharp decline in coverage observed in the 1:1,000 diluted samples enriched by NEB+MDA, while this drop was not observed in samples enriched by SWGA, which still held >95% coverage at 5X read depth. As observed with NEB+MDA, no sequencing errors were introduced during enrichment with SWGA. SWGA was subsequently used for enriching clinical specimens in this study.

### SWGA enrichment of clinical strains

SWGA on specimen EUHM-004 gave an average *T. pallidum* gDNA of 6.37 × 10^6^ ± 2.24 × 10^5^ copies/μl with 5.56% of the total DNA belonging to *T. pallidum* (Table 2). Next generation sequencing (NGS) on the MiSeq v2 (500 cycle) platform revealed 95.13% coverage across the *T. pallidum* genome at 5X read depth (Fig. 4; Table 2 and Fig. S2). Due to the overall low copy number obtained by standard DNA extraction, three specimens (EUHM-012, EUHM-013, and EUHM-014) were extracted using a large-scale method, yielding 31.5 ± 0.5, 122 ± 1.15, and 103 ± 6.55 *T. pallidum* gDNA copies/μl of extract, respectively (Table 1). For EUHM-012, we observed an average of 2.14 × 10^6^ ± 2.82 × 10^4^ *T. pallidum* gDNA copies/μl with 1.72% of the total DNA belonging to *T. pallidum* post-enrichment by SWGA (Table 2). Sequencing indicated a 93.98% coverage across the *T. pallidum* genome at 5X read depth (Fig. 4; Table 2 and Fig. S2).

When compared to EUHM-012, EUHM-013 had a higher *T. pallidum* gDNA copy number/μl at 5.16 × 10^6^ ± 2.20 × 10^5^ with 15.48% of the total DNA belonging to *T. pallidum* post-enrichment by SWGA (Table 2). The sequencing data correlated with the qPCR data, indicating a 98.56% coverage across the *T. pallidum* genome at 5X read depth (Fig. 4; Table 2 and Fig. S2). We also observed EUHM-014 held an increased *T. pallidum* gDNA copy number post-SWGA, with 2.57 × 10^6^ ± 2.21 × 10^5^copies/μl and 4.72% of the total DNA belonging to *T. pallidum* (Table 2). Upon sequencing, we observed 98.49% coverage across the *T. pallidum* genome at 5X read depth (Fig. 4; Table 2 and Fig. S2). The *T. pallidum* gDNA copy number for specimen STLC-001 was 7.42 × 10^6^ ± 7.20 × 10^5^ copies/μl with 8.34% of the total DNA belonging to *T. pallidum* (Table 2). The sequencing coverage was 95.94% at 5X read depth where 38.91% of the quality-controlled reads binned as *T. pallidum* along with a mean depth read coverage of 1,133.43X (Fig. 4; Table 2 and Fig. S2).

### Phylogenetic analysis and characterization of genotypic macrolide resistance

A whole genome phylogenetic tree was constructed using the genomes derived from the 5 clinical specimens and 2 isolates along with 126 high quality published *T. pallidum* genome sequences as of May 2021 (18–21, 29–31); see Table S5, methods in supplemental materials). Phylogenetic analysis revealed the presence of two dominant lineages (Nichols-like and Street-14 (SS14)-like), of which most strains belonged to the SS14-like lineage. We identified a total of four monophyletic clades within this phylogenetic tree with ≥ 30 bootstrap support (Fig. 6). Three of the clinically derived genomes from Atlanta, EUHM-004 (2019) EUHM-012 (2019), and EUHM-014 (2020), belonged to Nichols-like lineage (clade 1; n=12; Fig. 6). Interestingly, the other nine Nichols-like genomes in clade 1 were recent clinically derived genomes from Cuba (n=2; 2015-2016), Australia (n=1; 2014), France (n=2; 2012-2013) and United Kingdom (n=3; 2016) and together with the specimens from Atlanta, were distinct from the original Nichols strain isolated in 1912 and sent to different North American labs as *in vivo* derived clones. The three clinical specimens from Atlanta (EUHM-004, EUHM-012, and EUHM-014) and three clinically derived genomes from the United Kingdom isolated in 2016 (NL14, NL19 and NL17) carried the 23S rRNA A2058G mutation that confers macrolide resistance, suggesting a recent acquisition of this antibiotic resistance variant in the Nichols-like lineage.

**Fig 6.**
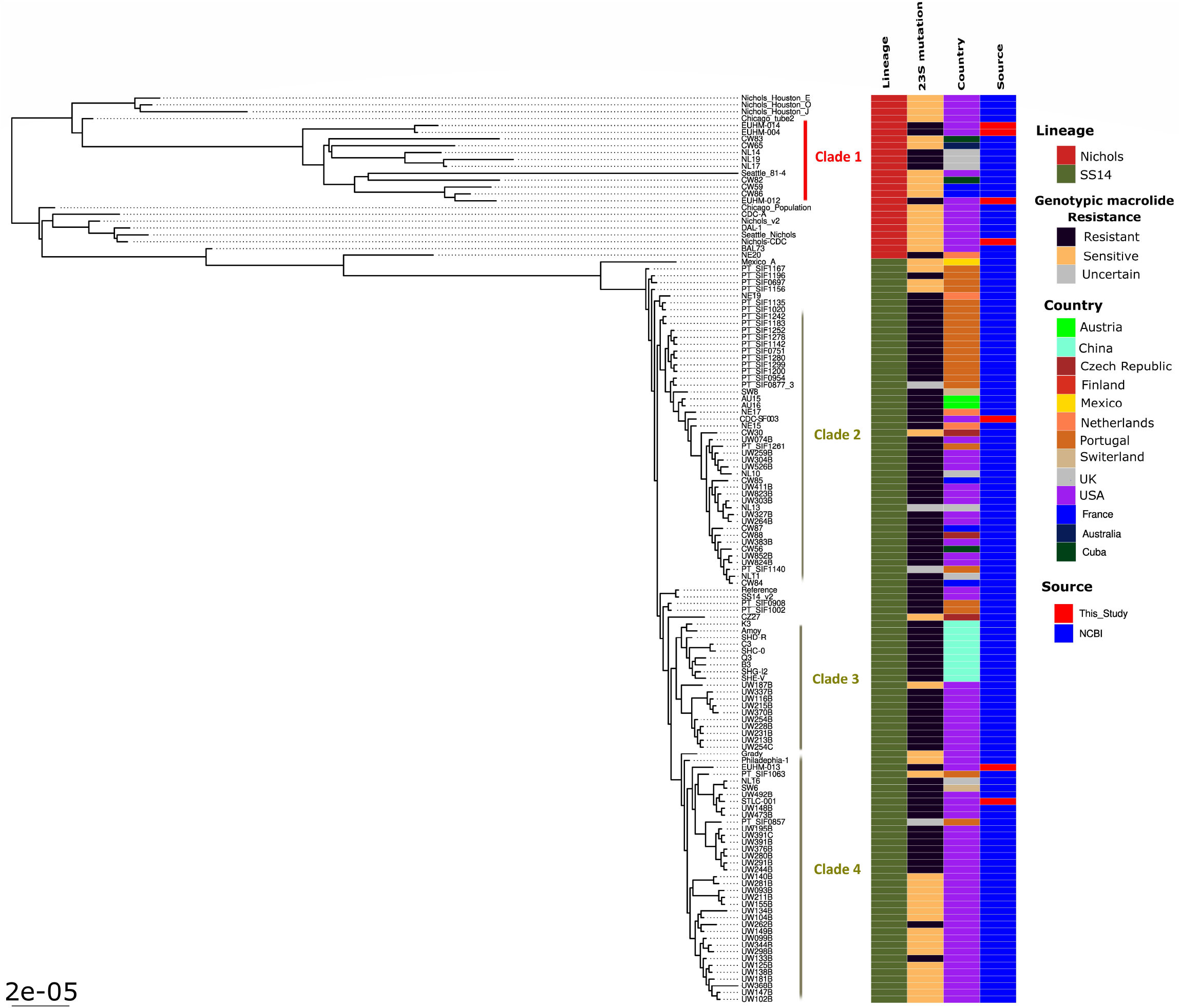
Maximum likelihood global phylogenetic tree of the 7 *T. pallidum* strains sequenced in this study along with 122 high quality (with 5x read depth covering >90% of the genome) publicly available *T. pallidum* genomes. The two major lineages, Nichols-like and SS14-like are highlighted along with presence of genotypic mutation responsible for macrolide resistance and country of origin.

Even though previous phylogenomic analyses indicated that SS14-lineage showed a polyphyletic structure, our phylogenetic analysis with a greater number of genomes showed the presence of 3 monophyletic clades (Clades 2, 3 and 4) (18, 20); Fig. 6. Clades 2 and 4 contained genomes clustered within the previously reported SS14Ω-A sub cluster, which also contained two clades corresponding to the clades 2 and 4 detected in this study, and contained genomes derived from Europe and North America; while clade 3 was like sub-cluster SS14Ω-B and composed of Chinese and North American derived *T. pallidum* genomes (18). The rabbit-derived isolate, CDC-SF003 (San Francisco, U.S; 2017) sequenced in this study, clustered within clade 2; while EUHM-013 (Atlanta, U.S; 2020) and STLC-001 (St. Louis, U.S; 2020) genomes clustered within clade 4. Sequence analysis showed that all 3 strains carried the A2058G antimicrobial resistance variant for macrolide resistance. Macrolide resistance strains were widespread among the SS14-lineage with higher proportion among the genomes in clades 2 and 3 compared to clade 4 genomes. The A2058G point mutation identified in 4 patient specimens and isolate CDC-SF003 was verified by real-time PCR testing of gDNA and SWGA-enriched samples (data not shown). There was inadequate sample for the fifth specimen to confirm the mutation by real-time PCR testing.

All the Nichols-like genomes derived from the NEB+MDA and SWGA 10-fold dilution series that contained *T. pallidum* reads mapped to ≥90% of the genome with at least 5X read depth formed a tight monophyletic clade (bootstrap support of 88/100) and clustered with the lab-derived Nichols-Houston-J genome (bootstrap support of 100/100), indicating that genomes generated from both methods are adequate to capture genetic variants required to perform a high-resolution phylogenetic analysis (Fig. S4).

## Discussion

WGS of *T. pallidum* is often challenging due to low bacterial loads or the difficulty of obtaining adequate samples for testing. In this study, we sought to develop a method for performing WGS from rabbit propagated isolates and clinical specimens containing lower *T. pallidum* numbers, leading us to investigate CpG capture and SWGA.

During the testing of the 10-fold dilution series of Nichols spiked samples, we observed increases in *T. pallidum* gDNA copy numbers and relative *T. pallidum* percent DNA in the neat to 1: 1,000 dilutions enriched by NEB+MDA when compared to the non-enriched inputs. There was no significant difference in the relative human RNPCt from dilution to dilution. While the neat to 1:100 dilution spiked samples demonstrated an average percent coverage exceeding 97.29% at 5X read depth across Nichols strain genome, genomic coverage for 1:1,000 to 1:1,0000 diluted spiked samples were poor. We observed that an input of >129 *T. pallidum* gDNA copies/μl can generate >95% coverage at 5X read depth from the Nichols strain post NEB+MDA enrichment. Post NEB+MDA enrichment of isolate CDC-SF003 demonstrated >98% coverage at 5X read depth across the *T. pallidum* genome. Variant analysis correlated with real-time PCR detection of the mutations associated with macrolide resistance in CDC-SF003. In addition, phylogenetics revealed that this strain belonged to the SS14 lineage, which correlated with its enhanced CDC typing method (ECDCT) strain type, 4d9f, as previously reported (11). While this enrichment method yielded good results with isolates, the fact that a majority of clinical specimens collected in this study had lower than 100 *T. pallidum* gDNA copies/μl extract led us to consider an alternative method.

We observed that samples enriched by SWGA using multiple primer sets exhibited a 10,000-fold increase in *T. pallidum* genome copy number, with Pal 9 and 11 producing the highest relative percent *T. pallidum* DNA at 29% and 31%, respectively. While we chose to work with Pal 11 as the optimal set, Pal 9 could also be a good alternative for enriching syphilis specimens. Further testing using Pal 11 showed that the limit of detection was increased when compared to the *T. pallidum* enrichment obtained with NEB+MDA, with significant increases in both *T. pallidum* gDNA copy number and percent *T. pallidum* across the 10-fold dilution series of spiked samples. Coverage across the *T. pallidum* genome exceeded 95% at 5X read depth for all diluted samples, apart from the 1:10,000 diluted samples. Interestingly, we observed that increasing the *T. pallidum* DNA input 100-fold resulted in a significant decrease in the presence of RNP post-enrichment. Our data shows that >14 *T. pallidum* gDNA copies/μl can generate at least 95% coverage at 5X read depth with the Nichols strain using the SWGA enrichment method, which translated well to the clinical specimens tested. While there was a decrease in coverage in one of the clinical specimens at 94.44% with 5X read depth when compared to the 98.62% coverage at 5X read depth observed in the 1:100 diluted Nichols isolates, this could be primarily due to the improved capabilities of the NovaSeq 6000 when compared to the MiSeq v2 (500 cycle) platform used for sequencing. Another possible reason for the variation in coverage could be due to the lower *T. pallidum* input copy number in the clinical specimens.

The genomes derived directly from the 5 clinical specimens using SWGA were phylogenetically associated with the representative lineages (either Nichols-like or SS14-like) and provided high levels of within lineage strain resolution, which is ideal for effective tracking of various strains circulating within a geographical area and outbreak investigations. In addition, the NGS methods described here can be used for macrolide resistance marker detection. As observed with NEB+MDA enrichment, azithromycin mutation detection performed on the SWGA enriched specimens matched the results obtained by real-time PCR, indicating that all clinical specimens contained the A2058G mutation. SWGA-based enrichment also enabled sequencing of specimens within the range of detection limits for real-time PCR assays, suggesting that our NGS workflow can be adapted for *T. pallidum* detection in metagenomic samples.

In terms of expense, both methods are cost-effective for enriching *T. pallidum* gDNA, and while SWGA is cheaper than NEB+MDA, sequencing reagents are the true limiting factor for WGS. With the recent advancements in large-scale sequencing platforms, overall sequencing costs can be further reduced. While NovaSeq 6000 has a much higher potential for multiplex sequencing, our data shows compatibility of these enrichments for both NovaSeq 6000 and MiSeq platforms.

While we successfully enriched *T. pallidum* whole genomes in clinical specimens, the success of SWGA is limited by the constraint on primer size, which may reduce the selectivity for the target genome. Phi29 functions best between 30-35°C, and ramp-down incubations have been shown as an effective means of utilizing larger primers with increased melting temperatures (26, 27, 32, 33). To help alleviate the constraints on primer size, we utilized a thermostable phi29 mutant which has a much higher optimal temperature at 45°C (34) compared to the 30-35°C functional range of the phi29 polymerase (26, 27). This higher optimal temperature permits the use of longer oligonucleotides to be used in the SWGA reaction, potentially increasing the selectivity for the *T. pallidum* genome. The phi29 mutant has also shown to be more efficient, with a 3hour exhaustion time when compared to the 8-16 hours required for the wild-type phi29 (34). In addition, the genome sequence data generated from NEB+MDA and SWGA enriched samples revealed that no sequencing errors were introduced during whole genome amplification methods.

Our results show that SWGA is more sensitive, less cumbersome, and a faster method for enriching clinical specimens when compared to NEB+MDA, allowing for WGS of *T. pallidum* with a minimum input of 14 gDNA copies/μl. In addition, the sequencing data generated is of sufficient quality to enable phylogenetic analyses and detection of mutations associated with azithromycin resistance. While NEB+MDA was unsuitable for the clinical specimens in this study, our data suggests that it can be used for DNA extracts containing >129 *T. pallidum* gDNA copies/μl.

## Materials and Methods

### Specimen collection, *T. pallidum* strains used for WGS, and real-time qPCR

Specimens used in this study were collected from men presenting with lesions of primary or secondary syphilis to the Emory Infectious Diseases Clinic, Emory University Hospital Midtown (EUHM) in Atlanta, GA and St. Louis County STD Clinic (STLC) in St. Louis, MO (Table 1). Patients were diagnosed with syphilis based on clinical presentation and serology testing. Fourteen swab specimens were collected in Aptima Multitest storage medium (Hologic, Inc., Marlborough, MA) at Emory Infectious Diseases Clinic and 1 specimen at St. Louis County STD Clinic (Table 1). All specimens were stored at −80°C until shipment on dry ice to the CDC. The *T. pallidum* Nichols reference strain was used for initial optimization and validation of the two enrichment methods. A recent rabbit propagated isolate, CDC-SF003, was also included for testing (Table 1; (11)). Prior to study commencement, local IRB approvals were obtained from Emory University and St. Louis County Department of Public Health, and the project was approved at CDC.

DNA was extracted from specimens and rabbit testis extracts using the QIAamp DNA Mini Kit (Qiagen, Germantown, MD) following the manufacturer’s recommendations. To increase our chances of successfully sequencing three specimens, large-scale DNA extraction was carried out on 1.5 ml of specimen using the QIAamp DNA Mini Kit with slight modifications for upscaling (Table 1). Briefly, proteinase K was added at 0.1X total sample volume. AL Buffer and ethanol were added at 1X total sample volume. Each sample was processed through a single column and eluted in 100 μl AE Buffer (Qiagen). DNA samples were tested by a real-time quantitative duplex PCR (qPCR) targeting the *polA* gene of *T. pallidum* and human RNase P gene (RNP) using a Rotor-Gene 6000 instrument (Qiagen) as previously described with modifications (11, 28); see additional methods in supplemental materials).

### NEBNext microbiome enrichment and multiple displacement amplification (MDA)

Initially, DNA concentration of extracts from clinical specimens and rabbit propagated strains were measured using the Qubit dsDNA HS assay (Thermo Fisher Scientific, Waltham, MA). Capture and removal of CpG methylated host DNA from samples were carried out using the NEBNext^®^ Microbiome DNA Enrichment Kit (NEB, New England Biolabs, Ipswich, MA), following the manufacturer’s recommendations with modifications (New England Biolabs, Ipswich, MA). For all samples tested, 250 ng of DNA was subjected to two rounds of bead capture using NEB and enriched treponemal gDNA was purified using AMPure XP beads (Beckman Coulter, Indianapolis, IN). Enriched DNA samples were stored at −20°C until MDA was performed. Multiple displacement amplification (MDA) was carried out using the REPLI-g Single Cell Kit following the manufacturer’s recommendations with slight modifications (Qiagen). MDA reactions were incubated at 30°C for 16 hr. Following amplification, the polymerase was inactivated at 65°C for 10 min, samples were purified with AMPure XP beads, and eluted with 100 μl 1X AE Buffer (Qiagen). For enrichment by MDA, no template controls were included to confirm the absence of *T. pallidum*.

A 10-fold dilution series on the Nichols strain was used to determine the limit of detection (LoD) for enrichment (see supplemental materials) with NEB+MDA followed by sequencing on an Illumina NovaSeq 6000. After DNA extraction, each dilution in the series was enriched by NEB+MDA, gDNA copy numbers estimated by *polA* qPCR, and sequencing performed in triplicate. Enriched samples were diluted 1:10 prior to measuring RNP amplification. The LoD was set at the minimum genome copy number required to generate a ≥5X read depth with ≥95% genome coverage compared to the reference genome.

### Selective whole genome amplification (SWGA) primer design, validation, and enrichment

Primers with an increased affinity to *T. pallidum* were identified using the *swga* Toolkit as previously described with slight modifications (https://www.github.com/eclarke/swga; (26); see supplemental materials). Eight primer sets (SWGA Pal 1 to 8), including 4 additional primer sets (SWGA Pal 9-12) generated by combining primers in the initial set (Table S1), were chosen for SWGA using the EquiPhi29 DNA Polymerase (Thermo Fisher Scientific, Waltham, MA). To account for the 3’-5’ exonuclease activity of the phi29 polymerase, all SWGA primers were generated with phosphorothioate bonds between the last two nucleotides at the 3’ end (Table S1). Each of the 12 primer sets were tested in triplicate against the spiked sample diluted to an estimated 100 *T. pallidum* gDNA copies/μl (see supplemental materials).

Prior to SWGA enrichment, samples were denatured for 5 min at 95°C by adding 2.5 μl of DNA to 2.5 μl reaction buffer, containing custom primers, then placed immediately on ice until the Equiphi29 master mix, prepared as per manufacturer’s recommendations, was added. Whole genome amplification was carried out following the manufacturer’s recommendations with modifications (Thermo Fisher Scientific; (34)). The reaction contained EquiPhi29 master mix, with EquiPhi29 Reaction Buffer at a final concentration of 1X, each primer at a final concentration of 4 μM, and nuclease-free H2O was added to a final reaction volume of 20 μl. Reaction tubes were gently mixed by pulse vortexing and incubated at 45°C for 3 hr. The reaction was stopped by inactivating the DNA polymerase at 65°C for 15 min. All reactions were purified using AMPure XP beads and eluted in 100 μl AE buffer (Qiagen). No template controls were included to confirm the absence of contaminate *T. pallidum* DNA.

Relative percent *T. pallidum* in each sample was calculated as shown in Figure S1. SWGA Pal 11 was chosen for testing the LoD for downstream genome sequencing post-SWGA enrichment using the 10-fold dilution series, excluding the undiluted (neat) spiked sample. All enriched samples were validated by *polA* real-time qPCR in triplicate.

### Sequencing and genome analysis of *T. pallidum* strains

Libraries were prepared using the NEBNext^®^ Ultra DNA Library Preparation Kit for NovaSeq and NEBNext^®^ Ultra II FS DNA Library Preparation Kit for MiSeq sequencing following the manufacturer’s recommendations (New England Biolabs, Ipswich, MA). For the validation experiments, sequencing was carried out on the Nichols reference strain using the Illumina NovaSeq 6000 platform following the manufacturer’s recommendations (Illumina, San Diego, CA). Sequencing of isolate CDC-SF003 and swab specimens were carried out using the MiSeq v2 (500 cycle) platform following the manufacturer’s recommendations (Illumina, San Diego, CA).

Sequenced genomic reads were first filtered from the raw sequencing dataset using Bowtie2 version 2.2.9 (35), which removed any contaminating human sequences using the h19 human reference genome and rabbit reference genome for rabbit propagated clinical isolates (36). Cutadapt version 1.8.3 was used to trim specified primers and adaptors, and to filter out reads below Phred quality scores of 15 and read length below 50 bps (37). PCR duplicates were removed using Clumpify (sourceforge.net/projects/bbmap/) with dedupe=t option to prevent biased coverage of genomic regions. *Treponemal* reads were selected using K-SLAM, a k-mer-based metagenomics taxonomic profiler, which used a database containing all bacterial and archaeal reference nucleotide sequences (38). The presence of *T. pallidum* sequences was also confirmed using Metaphlan2 (39). Prefiltered Treponemal reads were mapped against either the *T. pallidum* subsp. *pallidum* SS14 reference genome (*NC_021508.1*) or the *T. pallidum* subsp. *pallidum* Nichols reference genome (*NC_000919.1*) using BWA mem v2.12.0 (MapQ ≥ 20), followed by consensus sequence generation and estimation of sequencing depth and mapping statistics using samtools (options ‘depth’ and ‘mpileup’) and bcftools v1.9. The prefiltered *Treponemal* reads were also used to generate *de novo* short read assemblies using SPAdes 3.7.0 with the “careful” option (40–42).

To assess whether there were introductions of amplification-induced sequencing errors for both NEB and SWGA, we compared the sequencing data generated from all three replicates for each of the 10-fold dilution series for both protocols to the control genome generated directly from an unamplified DNA sample containing 10,000 gDNA copies/μl of *T. pallidum* Nichols. The same Nichols DNA was used for the dilution series and the unamplified sample. Essentially, the sequencing reads of the Nichols control genome were *denovo* assembled using SPAdes 3.7.0 as described above and the resultant contigs were arranged using abacas in reference to the *T. pallidum* Nichols reference genome (NC_021490.2) to generate a circular pseudo chromosome (43). All *T. pallidum* reads from all the NEB and SWGA 10-fold diluted samples were mapped against the control *T. pallidum* Nichols pseudo chromosome using Snippy v4.3.8 and checked for any genetic variants in the form of single nucleotide polymorphisms (SNPs) and insertions/deletions introduced during multiple amplification steps.

### Phylogenetic analyses

We used a reference mapping approach by mapping the filtered Treponemal reads to a custom versions of the *T. pallidum* SS14 reference genome (*NC_021508.1*) by masking around 30,000 nucleotide positions belonging to 12 repetitive *tpr* genes A-L, along with *arp* and TPANIC_0470 genes using bedtools v2.17.0 ‘maskfasta’ (44). Full length whole genome alignment were generated using Snippy v4.3.8(https://github.com/tseemann/snippy), which identifies variants using Freebayes v1.0.2 with a minimum 5X read coverage and 90% read concordance at a locus for each single nucleotide polymorphism (SNP) (45). Regions of increased density of homoplasious SNPs introduced by possible recombination events were predicted iteratively and masked using Gubbins (46). The final phylogenetic tree was reconstructed using RAxML on the recombination removed alignment using the GTR+GAMMA model of nucleotide substitution with a majority-rule consensus (MRE) convergence criterion, to reconstruct an ascertainment bias corrected (Stamatakis method) maximum-likelihood (ML) phylogeny (47).

Apart from the genomes sequenced in this study, 126 high quality (with at least 5x read depth covering >90% of the genome) *T. pallidum* genomes deposited in the NCBI’s Sequencing Read Archive (SRA) under BioProject number PRJEB20795 and PRJNA508872 were also included (18, 20). A second phylogenetic tree was also reconstructed by including all the genomes sequenced from the 10-fold dilution series for both NEB+MDA and SWGA enriched samples. Genomic sequencing data from samples included in the phylogenetic analyses covered at least 90% of the reference genome with 5X read depth. Variant calls for the A2058G and A2059G macrolide resistance mutations using genomic data were validated with a previously described real-time PCR assay (5).

### Statistical analyses

Statistical analyses were performed in R (R Foundation for Statistical Computing, Vienna, Austria) using the R companion software RStudio (Rstudio, Boston, MA). Statistical significance was determined by analysis of variance (ANOVA) and Tukey post hoc multiple comparisons tests. *T. pallidum* percent DNA were normalized through Log10 conversions. Quantitative data are presented as means ± standard error. Differences were considered statistically significant if a P < 0.05.

### Data availability

All sequencing data associated with this study were submitted to the National Center for Biotechnology Information’s sequence read archive (SRA) under the BioProject accession ID PRJNA744275.

## Acknowledgments

We thank Teressa Burns at the Emory University Hospital Midtown; Tamara Jones from the St Louis County STD Clinic; Yetty Fakile, Kevin Pettus, and Jack Cartee at CDC’s Division of STD Prevention; The veterinary staff at the CDC’s Comparative Medicine Branch; Mark Itsko at CDC’s Division of Bacterial Diseases; and Nikhat Sulaiman and Justin Lee at CDC’s Division of Scientific Resources for their assistance, consults, and support throughout this study. This work was made possible through CDC’s Division of STD Prevention with support from the Advanced Molecular Detection (AMD) program, and in part by an appointment (CMT) to the Research Participation Program at the CDC, administered by the Oak Ridge Institute for Science and Education through an interagency agreement between the US Department of Energy and CDC.

## Author Contributions

Allan Pillay and Ellen N. Kersh conceived the study. Allan Pillay, Charles M. Thurlow, Cheng Chen, and Lilia Ganova-Raeva designed the study. Charles M. Thurlow and Allan Pillay designed the enrichment protocols. Charles M. Thurlow designed the SWGA specific custom primer sets used during this study and performed all enrichment experiments. Charles M. Thurlow, Allan Pillay, Samantha S. Katz, Lara Pereira, Alyssa Debra, Kendra Vilfort, Yongcheng Sun, Kai-Hua Chi, and Damien Danavall performed the laboratory experiments and assisted with specimen collection. Kimberly Workowski, Stephanie E. Cohen, Hilary Reno, and Susan S. Philip collected clinical specimens and patient data. Mark Burroughs, Mili Sheth, and Charles M. Thurlow performed Illumina sequencing. Sandeep J. Joseph performed the bioinformatic analyses of the genomic data, phylogenetic analysis and contributed to the generation of tables and figures. Charles M. Thurlow and Sandeep J. Joseph performed data analysis. Charles M. Thurlow wrote and prepared the manuscript with oversight by Allan Pillay and contributions from Sandeep J. Joseph and Weiping Cao, which was reviewed by all authors for revisions.

## Disclaimer

The findings and conclusions in this report are those of the authors and do not necessarily represent the official position of the Centers for Disease Control and Prevention. We declare that there are no competing interests.

