## Supplemental Materials & Methods for "Selective whole genome amplification as a tool to enrich specimens with low *Treponema pallidum* genomic DNA copies for whole genome sequencing"

### Supplemental Materials and Methods

**Real-time qPCR.** All standard extracted samples were tested by quantitative PCR (qPCR) in triplicate apart from specimens extracted by large-scale processing, which were tested in duplicate to preserve DNA. Each replicate reaction consisted of 10 µl DNA; *polA* (TP-polA-FP, TP-polA-RP) and RNP (RNP-FP and RNP-RP) primers at a final concentration of 300 nM and 80 nM, respectively; FAM (TP-FAM23) and Quasar 670 (RNP-Q670) TaqMan probes at a final concentration of 200 nM and 80 nM, respectively; and 25 µl of PerfectA qPCR Supermix (Quanta Biosciences, Beverly, MA) in a 50 µl reaction. All primers and probes were synthesized at the Oligo Synthesis Laboratory in the CDC's Division of Scientific Resources (Table S2). All test samples were run alongside *T. pallidum* and human DNA positive controls, and no template controls (NTC). $RNP_{Ct}$ was calculated by aligning the RNP amplification against a predetermined standard curve generated using a 10-fold serial dilution of human genomic DNA (gDNA; Millipore Sigma, Burlington, MA). Standard curve generation and quantification were carried out as per the Rotorgene 6000 manufacturer's recommendations (Qiagen, Germantown, MD). Standardization of the relative $polA_{Ct}$ in each sample was carried out similarly to the $RNP_{Ct}$ standardization. The *polA* standard curve was constructed by using DNA quantified from the *T. pallidum* Nichols isolate using relative dark-field microscopy counts as previously described (1). The test samples were also run alongside *T. pallidum* Nichols samples with known *polA* copies for quantifying the relative gDNA copies/µl for each sample. Genomic DNA was measured using the Qubit dsDNA BR/HS kit(s), and relative *T. pallidum* percent in all samples were calculated using both the *polA* copies/µl and DNA concentrations in the equations in supplemental Fig. S5.

#### ***T. pallidum*** **Nichols spiked samples for determining the enrichment limit of detection (LoD)**

For the spiked samples used for determining the LoD of each enrichment, we generated a 10-fold dilution series of *T. pallidum* Nichols and human DNA in Aptima Multitest Transport Medium (Hologic, Inc., Marlborough, MA). We targeted 10,000 copies/µl to 1 copy/µl of *T. pallidum* DNA based on darkfield microscopy counts as described above. $RNP_{Ct}$ values of the standard extracted clinical specimens were used when determining the concentration of the purified human gDNA (Millipore Sigma) spiked into mock samples. DNA was extracted from the diluted samples as described above, and *T. pallidum polA* qPCR was performed to verify the genome copy number. The standardized $RNP_{Ct}$ was also measured by real-time PCR to confirm that the human gDNA concentration was comparable amongst all dilutions in the series. Triplicate reactions of each sample in the dilution series were validated against *polA* and RNP standard curves.

**SWGA primer design**

*T. pallidum* Nichols *(NC_000919.1)* was used as the target genome and the *Homo sapiens* genome (assembly GRCh38.p12) was used as the background genome. After importing the Nichols and human genomes, the command *swga count* was used to count kmers in the foreground that range between 5-12 bp in length. Stringency was placed on primers counted so that the minimum foreground binding distribution ranged from 11-100 binding sites with a maximum background binding distribution ranging from 10,000-22,000 binding sites. The counted kmers were further separated using *swga filter* using the previous *swga count* parameters. The s*wga find_sets* command was used to call primer sets with a range of 2-40 primers and a minimum background primer binding distance between 60,000-100,000 bp. Primer sets Pal 1-Pal 8 were chosen for additional testing by MDA (Table S2) based on their calculated binding distributions, Gini coefficients on binding distributions, and set scores (Table S3). Primers used for primer sets Pal 9-12, were derived from primer sets Pal 1- Pal 8 based on the above binding distributions (Table S2).

**Supplementary Tables**

**Table S1:** NEB/SWGA dilution stats

**Table S2:** List of Primers and Probes used in this study

| **Primer Set ID** | **Primer Name** | **Sequence (5' to 3')** | **References** |
|---|---|---|---|
| ***T. pallidum polA*** | TP-*polA*-FP | CAGGATCCGGCATATGTCC | Pereira *et al.* 2020 |
| | TP-*polA*-RP | AAGTGTGAGCGTCTCATCATTCC | Pereira *et al.* 2020 |
| | TP-*polA*-Probe | *FAM*-CTGTCATGCACCAGCTTCGACGTCTT-*BHQ1* | Pereira *et al.* 2020 |
| **Human RNP** | RNP-FP | CCAAGTGTGAGGGCTGAAAAG | This study |
| | RNP-RP | TGTTGTGGCTGATGAACTATAAAAGG | This study |
| | RNP-probe | *QUAS670*-CCCCAGTCTCTGTCAGCACTCCCTTC-*BHQ3* | This study |
| **SWGA-Pal 1** | SWGA-Pal 1 | TTGCG*C*G | This study |
| **SWGA-Pal 2** | SWGA-Pal 2 | CGCGC*A*A | This study |
| **SWGA-Pal 3** | SWGA-Pal 3.1 | AACTTT*T*T | This study |
| | SWGA-Pal 3.2 | CGGTAGT*A*G | This study |
| | SWGA-Pal 3.3 | TACGGAT*A*G | This study |
| | SWGA-Pal 3.4 | TCGTATA*C*G | This study |
| | SWGA-Pal 3.5 | TTTTTGAA*C*G | This study |
| **SWGA-Pal 4** | SWGA-Pal 4.1 | CGCGA*A*A | This study |
| | SWGA-Pal 4.2 | CGTAC*C*G | This study |
| | SWGA-Pal 4.3 | CGTAC*G*A | This study |
| | SWGA-Pal 4.4 | CGTAT*C*G | This study |
| | SWGA-Pal 4.5 | TACGC*G*T | This study |
| **SWGA-Pal 5** | SWGA-Pal 4.1 | CGCGA*A*A | This study |
| | SWGA-Pal 4.2 | CGTAC*C*G | This study |
| | SWGA-Pal 4.4 | CGTAT*C*G | This study |
| | SWGA-Pal 4.5 | TACGC*G*T | This study |
| | SWGA-Pal 5.1 | CGCGT*A*A | This study |
| **SWGA-Pal 6** | SWGA-Pal 6.1 | AACGTAT*C*G | This study |
| | SWGA-Pal 6.2 | CGCGTA*A*A | This study |
| | SWGA-Pal 6.3 | CGCGTAT*T*A | This study |
| | SWGA-Pal 6.4 | CGCTAT*C*G | This study |
| | SWGA-Pal 6.5 | CGGTAAAA*A*A | This study |
| | SWGA-Pal 6.6 | CGGTAT*C*G | This study |
| | SWGA-Pal 6.7 | CGTAAAA*C*G | This study |
| | SWGA-Pal 6.8 | CGTATC*G*G | This study |
| | SWGA-Pal 6.9 | CGTATCG*T*A | This study |
| | SWGA-Pal 6.10 | CGTATTC*G*T | This study |
| | SWGA-Pal 6.11 | GTTTCGT*A*C | This study |

| | | | |
|---|---|---|---|
| | SWGA-Pal 6.12 | TCGTAA*C*G | This study |
| | SWGA-Pal 6.13 | TCGTAT*C*G | This study |
| | SWGA-Pal 6.14 | TCGTATT*C*G | This study |
| | SWGA-Pal 6.15 | ATATCGT*C*G | This study |
| **SWGA-Pal 7** | SWGA-Pal 7.1 | AACGATA*C*G | This study |
| | SWGA-Pal 7.2 | ACCGATA*G*T | This study |
| | SWGA-Pal 7.3 | ATCGATA*C*G | This study |
| | SWGA-Pal 7.4 | CCGATA*C*G | This study |
| | SWGA-Pal 7.5 | CGAATAC*G*A | This study |
| | SWGA-Pal 7.6 | CGACGAT*A*T | This study |
| | SWGA-Pal 7.7 | CGATAAA*C*G | This study |
| | SWGA-Pal 7.8 | CGATAG*C*G | This study |
| | SWGA-Pal 7.9 | CGCGAT*T*A | This study |
| | SWGA-Pal 7.10 | CGTCGAT*A*A | This study |
| | SWGA-Pal 7.11 | GCGAATA*A*C | This study |
| | SWGA-Pal 7.12 | GTACGAA*A*C | This study |
| | SWGA-Pal 7.13 | TACCGGA*T*A | This study |
| | SWGA-Pal 7.14 | TCGATAA*C*C | This study |
| | SWGA-Pal 6.5 | CGGTAAAA*A*A | This study |
| **SWGA-Pal 8** | SWGA-Pal 6.1 | AACGTAT*C*G | This study |
| | SWGA-Pal 8.1 | ACTATCG*G*T | This study |
| | SWGA-Pal 6.2 | CGCGTA*A*A | This study |
| | SWGA-Pal 6.3 | CGCGTAT*T*A | This study |
| | SWGA-Pal 6.4 | CGCTAT*C*G | This study |
| | SWGA-Pal 6.5 | CGGTAAAA*A*A | This study |
| | SWGA-Pal 6.6 | CGGTAT*C*G | This study |
| | SWGA-Pal 6.7 | CGTAAAA*C*G | This study |
| | SWGA-Pal 6.8 | CGTAAC*G*G | This study |
| | SWGA-Pal 8.2 | CGTATC*G*G | This study |
| | SWGA-Pal 6.9 | CGTATCG*T*A | This study |
| | SWGA-Pal 6.10 | CGTATTC*G*T | This study |
| | SWGA-Pal 8.3 | GTCGTAT*C*C | This study |
| | SWGA-Pal 8.4 | GTTATTC*G*C | This study |
| | SWGA-Pal 6.11 | GTTTCGT*A*C | This study |
| | SWGA-Pal 8.5 | TATACTC*G*C | This study |
| | SWGA-Pal 8.6 | TATCCGG*T*A | This study |
| | SWGA-Pal 8.7 | TATCGT*C*G | This study |
| | SWGA-Pal 6.12 | TCGTAA*C*G | This study |
| | SWGA-Pal 6.13 | TCGTAT*C*G | This study |
| | SWGA-Pal 6.14 | TCGTATT*C*G | This study |
| **SWGA-Pal 9** | SWGA-Pal 4.1 | CGCGA*A*A | This study |
| | SWGA-Pal 4.2 | CGTAC*C*G | This study |
| | SWGA-Pal 4.3 | CGTAC*G*A | This study |
| | SWGA-Pal 4.4 | CGTAT*C*G | This study |
| | SWGA-Pal 4.5 | TACGC*G*T | This study |
| | SWGA-Pal 5.1 | CGCGT*A*A | This study |
| **SWGA-Pal 10** | SWGA-Pal 4.1 | CGCGA*A*A | This study |
| | SWGA-Pal 4.2 | CGTAC*C*G | This study |
| | SWGA-Pal 4.3 | CGTAC*G*A | This study |

| | | | |
|---|---|---|---|
| | SWGA-Pal 4.4 | CGTAT*C*G | This study |
| | SWGA-Pal 4.5 | TACGC*G*T | This study |
| | SWGA-Pal 2 | CGCGC*A*A | This study |
| **SWGA-Pal 11** | SWGA-Pal 4.1 | CGCGA*A*A | This study |
| | SWGA-Pal 4.2 | CGTAC*C*G | This study |
| | SWGA-Pal 4.4 | CGTAT*C*G | This study |
| | SWGA-Pal 4.5 | TACGC*G*T | This study |
| | SWGA-Pal 5.1 | CGCGT*A*A | This study |
| | SWGA-Pal 2 | CGCGC*A*A | This study |
| **SWGA-Pal 12** | SWGA-Pal 4.1 | CGCGA*A*A | This study |
| | SWGA-Pal 4.2 | CGTAC*C*G | This study |
| | SWGA-Pal 4.3 | CGTAC*G*A | This study |
| | SWGA-Pal 4.4 | CGTAT*C*G | This study |
| | SWGA-Pal 4.5 | TACGC*G*T | This study |
| | SWGA-Pal 5.1 | CGCGT*A*A | This study |
| | SWGA-Pal 2 | CGCGC*A*A | This study |

[*]Nucleotides with phosphorothioate bonds; FAM: 6-carboxyfluorescein; QUAS670: Quasar 670; BHQ: Black hole quencher

**Table S3.** Binding Distribution of the *swga* Toolkit generated Primer Sets to the *Treponema pallidum* Genome and the *Homo Sapiens* Genome.

| SWGA Set ID | *T. pallidum* Distribution Mean[a] | *H. Sapiens* Distribution Mean[a] | *T. pallidum* Distribution Gini[a] | SWGA Set Score[a] |
|---|---|---|---|---|
| **SWGA-Pal 1** | 1,768.95 | 433,540.81 | 0.52 | 0.002 |
| **SWGA-Pal 2** | 1,768.95 | 433,540.81 | 0.52 | 0.002 |
| **SWGA-Pal 3** | 9,573.16 | 643,287.07 | 0.49 | 0.007 |
| **SWGA-Pal 4** | 1,023.54 | 113,241.19 | 0.53 | 0.004 |
| **SWGA-Pal 5** | 986.32 | 114,397.60 | 0.56 | 0.004 |
| **SWGA-Pal 6** | 2,982.20 | 243,020.67 | 0.53 | 0.006 |
| **SWGA-Pal 7** | 4,039.72 | 229,188.07 | 0.45 | 0.008 |
| **SWGA-Pal 8** | 2,383.27 | 147,921.14 | 0.52 | 0.008 |

[a]*T. pallidum* Distribution Mean, Homo Sapiens Distribution Mean, *T. pallidum* Gini, and SWGA Set Score were all calculated using the swga Toolkit.

**Table S4:** SWGA primer validation stats

**Table S5:** *T. pallidum* strains included in the phylogenetic tree

**Supplementary Figures**

**Fig S1.** Coverage plots for the *T. pallidum* Nichols undiluted and 10-fold dilution series enriched using the NEB+MDA. Quality controlled *T. pallidum* reads were mapped to the Nichols reference genome (*NC_000919.1*) and the $\log_{10}$ of the coverage at each mapped nucleotide position is shown on the x-axis**.** Samples were sequenced using the Illumina NovaSeq 6000 platform.

**Fig S2.** Coverage plots for the *T. pallidum* Nichols, Emory and St. Louis derived clinical specimens, and clinical isolate CDC-SF003. *T. pallidum* Nichols reference isolate quality-controlled *T. pallidum* reads were mapped to the Nichols reference genome (*NC_000919.1).* An input number of 1,000 genomic copies/µl *T. pallidum* was used for the Nichols non-enriched and SWGA enriched samples. Quality controlled *T. pallidum* reads for the Emory and St. Louis derived specimens and CDC-SF003 were mapped to the SS14 reference genome *(NC_021508.1*). The log$_{10}$ of the coverage at each mapped nucleotide position is shown on the x-axis. All samples were sequenced using the Illumina MiSeq v2 (500 cycle) platform.

**Fig S3.** Coverage plots for the *T. pallidum* Nichols dilution series by SWGA. Plots range from the lowest concentration *T. pallidum* Nichols mock samples (1:10,000) to the highest concentration (1:10) enriched using SWGA. Quality controlled *T. pallidum* reads were mapped to the Nichols reference genome (*NC_000919.1*) and the log$_{10}$ of the coverage at each mapped nucleotide position is shown on the x-axis**.** Samples were sequenced using the Illumina NovaSeq 6000 platform.

**Fig S4**. Maximum likelihood global phylogenetic tree with all the genomes sequenced from all the enriched Nichols dilution series with NEB +MDA and SWGA that generated at least 5X read depth covering at least 90% of the genome. All highlighted samples were sequenced using the Illumina NovaSeq 6000 platform.

**Fig S5**. *T. pallidum* DNA Ratio and Percent *T. pallidum* DNA equations. *T. pallidum* copy number was calculated from the *polA* copies/µL extract. The DNA concentration was measured using the Qubit dsDNA HS assay.
