## Supplementary figures and images for "Selective whole genome amplification as a tool to enrich specimens with low *Treponema pallidum* genomic DNA copies for whole genome sequencing"

### Supplementary Figure S1

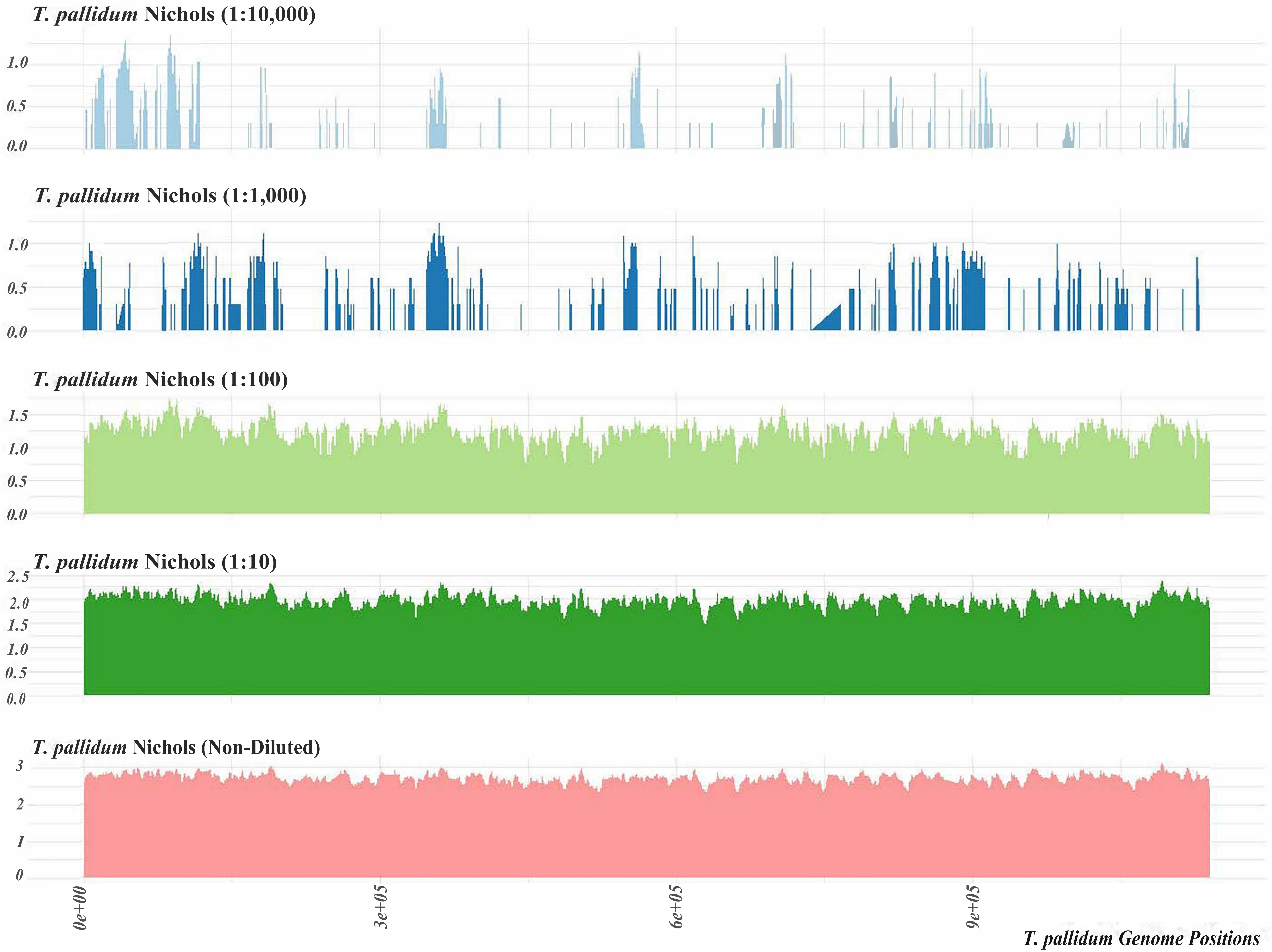

### Supplementary Figure S2

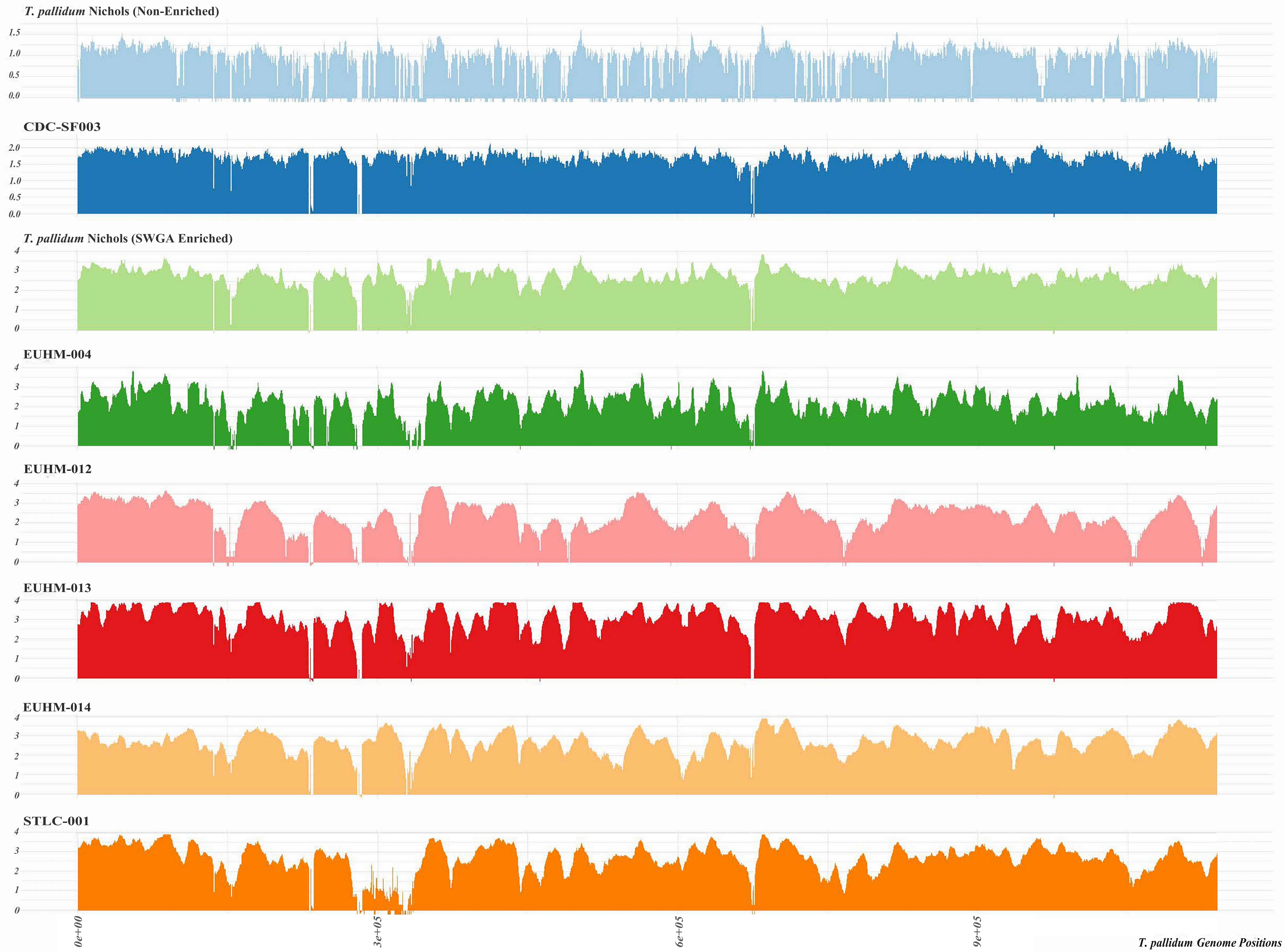

### Supplementary Figure S3

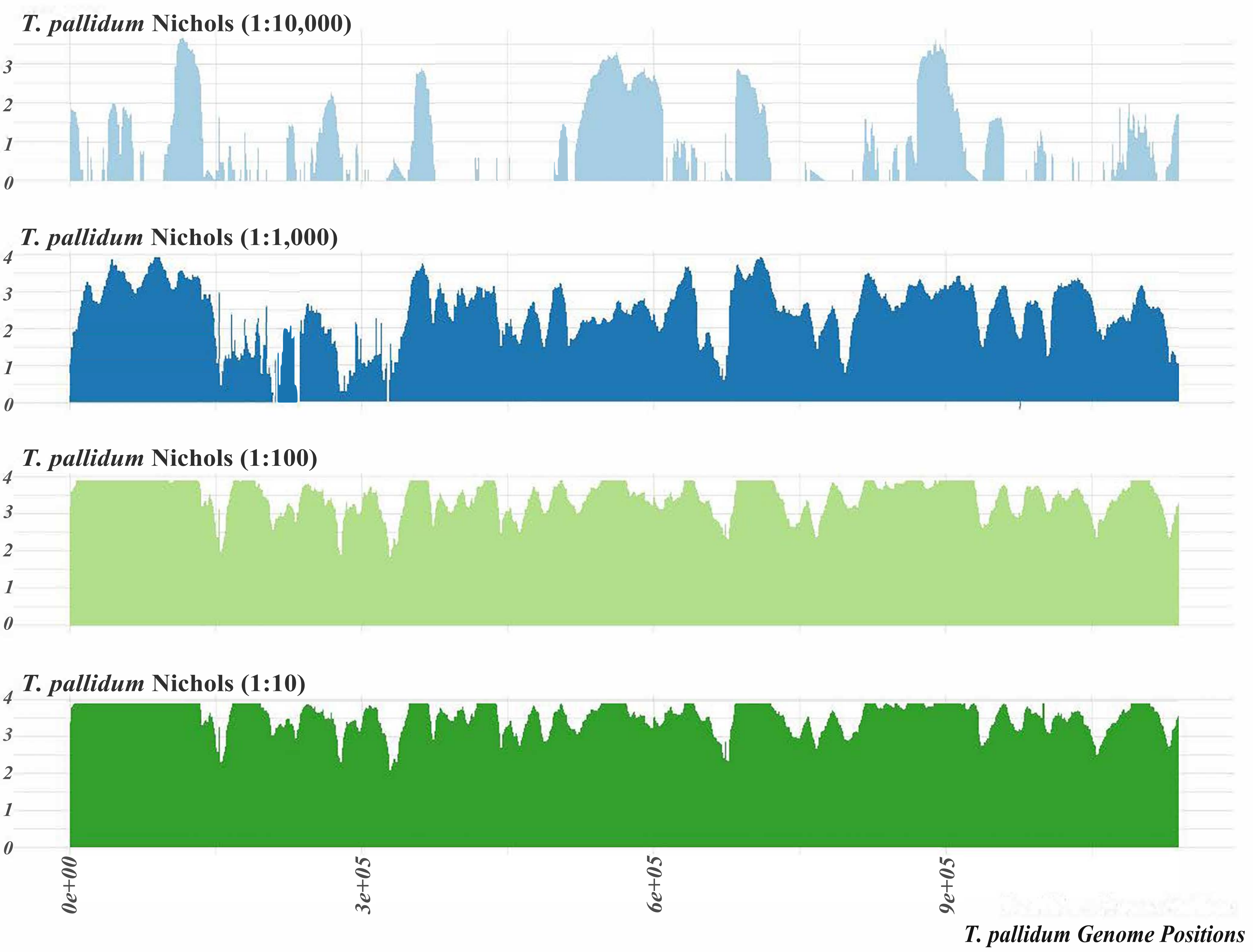

### Supplementary Figure S4

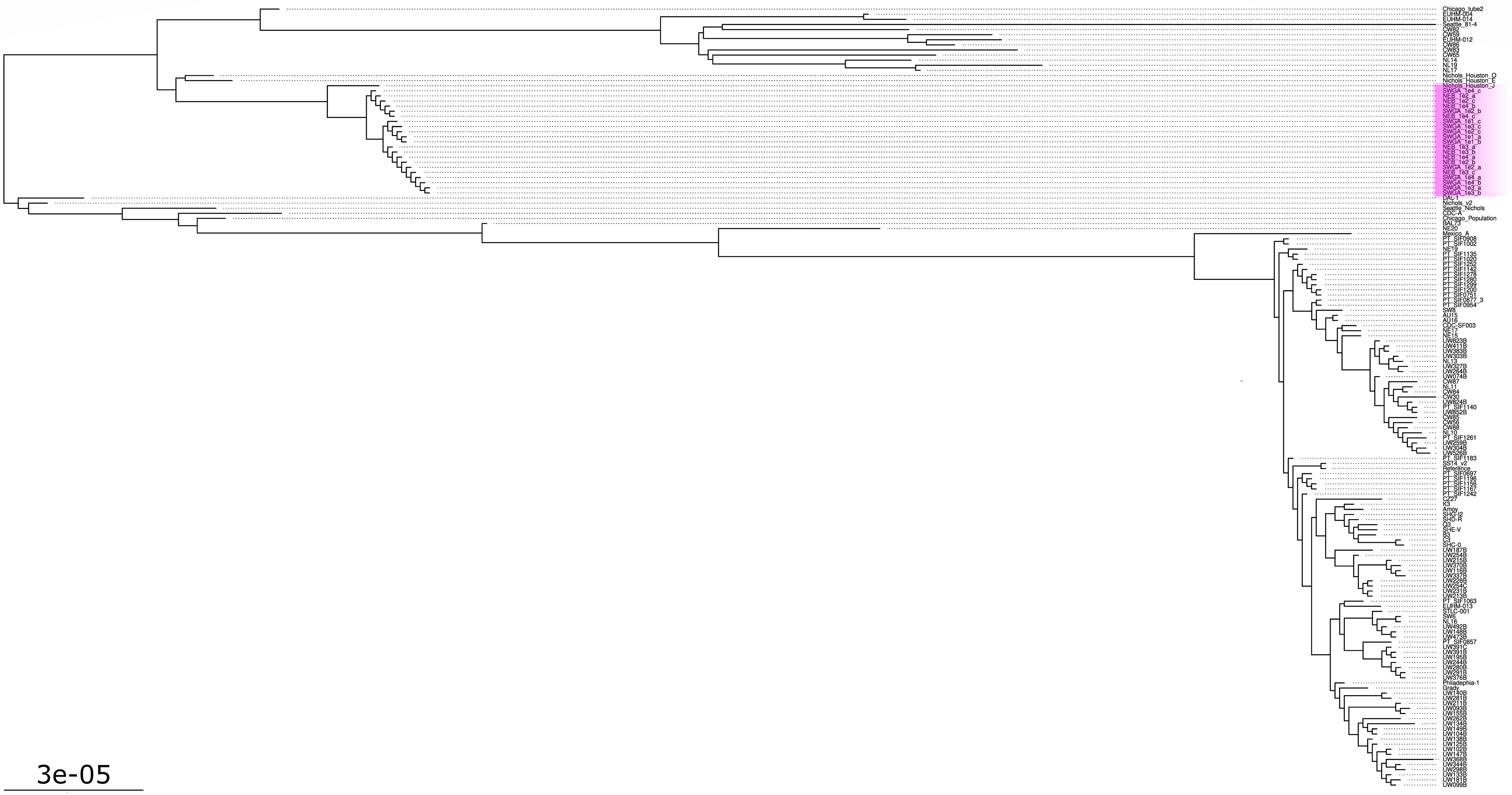
