## Supplementary Figure S5 for "Selective whole genome amplification as a tool to enrich specimens with low *Treponema pallidum* genomic DNA copies for whole genome sequencing"

**For Calculating Concentration of *T. pallidum* in sample (based on qPCR copy number):**

$$T.pallidum\ DNA\ Mass\ (ng) = \left(\frac{801757.42}{Calculated\ T.pallidum\ Copy\ Number}\right)\ x\ Elution\ Volume$$

** 801757.42 is the calculated copies of T. pallidum needed for 1 ng of DNA.*

**For Calculating DNA Ratio (*T. pallidum* DNA: total DNA):**

$$T.pallidum\ DNA\ Ratio = \left(\frac{Total\ DNA\ Concentration\ \left(\frac{ng}{\mu L}\right) x\ Elution\ Volume}{T.pallidum\ DNA\ Mass\ (ng)}\right)$$

**For Calculating Percent of Total DNA Belonging to *T. pallidum*:**

$$T.pallidum\ DNA\ ratio\ (percent) = \left(\frac{1}{T.pallidum\ DNA\ Ratio}\right)\ x\ 100$$
