## Supplementary Table S1, S4, S5 for "Selective whole genome amplification as a tool to enrich specimens with low *Treponema pallidum* genomic DNA copies for whole genome sequencing"

| ID | Enrichment method* | *T. pallidum* genome copies/µl in spiked samples (pre-enrichment) | $RNP_{Ct}$ in spiked samples (pre-enrichment) | *T. pallidum* genome copies/µl in spiked samples (post-enrichment) | $RNP_{Ct}$ in spiked samples (post-enrichment) | # of Raw read pairs | Total # of read pairs after human removal, adaptor triming and quality assessment | # of read pairs classified as T. pallidum | % of the read pairs classified as *T. pallidum* | Mean Read Depth | % of genome covered >=1X | % of genome covered >=5X | % of genome covered >=10X |
|---|---|---|---|---|---|---|---|---|---|---|---|---|---|
| NEB+MDA_1:10,000_rep1 | NEB+MDA | 1.50 | 24.95 | 726.00 | 30.81 | 22,987,457.00 | 12,847,858.00 | 1,643.00 | 0.01 | 0.67 | 20.27 | 4.98 | 0.93 |
| NEB+MDA_1:10,000_rep2 | NEB+MDA | 1.40 | 24.98 | 110.00 | 31.16 | 21,168,862.00 | 11,669,225.00 | 1,266.00 | 0.01 | 0.51 | 14.95 | 3.64 | 0.87 |
| NEB+MDA_1:10,000_rep3 | NEB+MDA | 1.80 | 25.10 | 2,056.00 | 32.29 | 18,995,197.00 | 10,345,644.00 | 1,885.00 | 0.02 | 0.76 | 22.69 | 5.78 | 0.85 |
| NEB+MDA_1:1,000_rep1 | NEB+MDA | 12.90 | 25.00 | 11,871.00 | 29.99 | 28,397,459.00 | 13,544,664.00 | 43,634.00 | 0.32 | 18.17 | 79.46 | 64.28 | 49.02 |
| NEB+MDA_1:1,000_rep2 | NEB+MDA | 13.00 | 25.07 | 3,649.00 | 31.22 | 16,048,959.00 | 9,009,594.00 | 3,160.00 | 0.04 | 1.28 | 36.68 | 9.89 | 1.12 |
| NEB+MDA_1:1,000_rep3 | NEB+MDA | 13.80 | 25.13 | 10,448.00 | 32.03 | 22,455,122.00 | 12,693,326.00 | 2,481.00 | 0.02 | 1.00 | 31.91 | 7.08 | 0.80 |
| NEB+MDA_1:100_rep1 | NEB+MDA | 138.90 | 24.84 | 148,316.00 | 31.30 | 12,130,073.00 | 7,325,455.00 | 33,199.00 | 0.45 | 13.77 | 99.93 | 95.18 | 71.49 |
| NEB+MDA_1:100_rep2 | NEB+MDA | 111.80 | 25.10 | 131,899.00 | 29.38 | 23,117,357.00 | 15,576,108.00 | 93,578.00 | 0.60 | 39.39 | 99.99 | 99.90 | 99.34 |
| NEB+MDA_1:100_rep3 | NEB+MDA | 137.60 | 24.82 | 104,883.00 | 30.06 | 21,990,019.00 | 11,031,554.00 | 39,001.00 | 0.35 | 16.48 | 99.73 | 96.81 | 81.60 |
| NEB+MDA_1:10_rep1 | NEB+MDA | 1,005.70 | 25.02 | 779,820.00 | 30.25 | 22,900,903.00 | 11,355,458.00 | 197,512.00 | 1.74 | 83.37 | 99.99 | 99.98 | 99.97 |
| NEB+MDA_1:10_rep2 | NEB+MDA | 1,035.40 | 25.12 | 852,750.00 | 31.41 | 22,840,971.00 | 11,365,682.00 | 212,416.00 | 1.87 | 89.72 | 99.99 | 99.99 | 99.98 |
| NEB+MDA_1:10_rep3 | NEB+MDA | 1,008.90 | 25.03 | 721,630.00 | 31.79 | 20,088,396.00 | 9,951,961.00 | 212,200.00 | 2.13 | 89.52 | 99.99 | 99.99 | 99.98 |
| NEB+MDA_neat_rep1 | NEB+MDA | 10,497.30 | 25.15 | 7,216,300.00 | 29.70 | 17,256,708.00 | 11,262,434.00 | 1,185,242.00 | 10.52 | 501.75 | 99.99 | 99.99 | 99.99 |
| NEB+MDA_neat_rep2 | NEB+MDA | 10,955.50 | 25.03 | 6,468,500.00 | 30.88 | 17,065,993.00 | 11,392,200.00 | 1,096,610.00 | 9.63 | 464.28 | 99.99 | 99.99 | 99.99 |
| NEB+MDA_neat_rep3 | NEB+MDA | 11,745.70 | 25.03 | 6,334,100.00 | 27.26 | 15,369,076.00 | 10,138,586.00 | 889,993.00 | 8.78 | 375.86 | 99.99 | 99.99 | 99.99 |
| SWGA_1:10,000_rep1 | SWGA | 1.50 | 24.95 | 2,438,800.00 | 28.67 | 23,192,918.00 | 15,017,442.00 | 393,396.00 | 2.62 | 164.85 | 49.68 | 34.98 | 29.47 |
| SWGA_1:10,000_rep2 | SWGA | 1.40 | 24.98 | 363,200.00 | 28.44 | 23,266,163.00 | 14,875,744.00 | 157,531.00 | 1.06 | 65.82 | 52.39 | 37.10 | 31.86 |
| SWGA_1:10,000_rep3 | SWGA | 1.80 | 25.10 | 513,900.00 | 28.85 | 27,225,312.00 | 16,590,056.00 | 162,696.00 | 0.98 | 68.03 | 61.20 | 43.30 | 36.01 |
| SWGA_1:1,000_rep1 | SWGA | 12.90 | 25.00 | 3,692,900.00 | 28.58 | 24,424,928.00 | 15,208,623.00 | 3,738,326.00 | 24.58 | 799.43 | 98.37 | 93.97 | 89.49 |
| SWGA_1:1,000_rep2 | SWGA | 13.00 | 25.07 | 5,512,300.00 | 28.88 | 28,291,268.00 | 18,868,094.00 | 5,334,480.00 | 28.27 | 2,067.50 | 99.75 | 98.87 | 98.35 |
| SWGA_1:1,000_rep3 | SWGA | 13.80 | 25.13 | 4,564,500.00 | 28.90 | 30,982,443.00 | 22,497,319.00 | 4,697,593.00 | 20.88 | 1,746.35 | 98.59 | 93.50 | 88.94 |
| SWGA_1:100_rep1 | SWGA | 138.90 | 24.84 | 14,364,100.00 | 28.94 | 31,426,441.00 | 25,124,204.00 | 14,672,298.00 | 58.40 | 4,123.81 | 99.99 | 99.99 | 99.98 |
| SWGA_1:100_rep2 | SWGA | 111.80 | 25.10 | 18,046,200.00 | 29.13 | 30,783,339.00 | 24,372,409.00 | 14,780,695.00 | 60.65 | 3,997.63 | 99.99 | 99.98 | 99.97 |
| SWGA_1:100_rep3 | SWGA | 137.60 | 24.82 | 13,604,500.00 | 29.28 | 30,984,647.00 | 25,453,153.00 | 13,520,963.00 | 53.12 | 4,161.87 | 99.99 | 99.99 | 99.98 |
| SWGA_1:10_rep1 | SWGA | 1,005.70 | 25.02 | 22,526,900.00 | 29.29 | 23,096,970.00 | 19,307,311.00 | 15,069,745.00 | 78.05 | 4,519.92 | 99.99 | 99.99 | 99.99 |
| SWGA_1:10_rep2 | SWGA | 1,035.40 | 25.12 | 18,365,400.00 | 29.30 | 26,561,373.00 | 22,049,992.00 | 16,534,836.00 | 74.99 | 4,705.41 | 99.99 | 99.99 | 99.99 |
| SWGA_1:10_rep3 | SWGA | 1,008.90 | 25.03 | 20,287,300.00 | 29.50 | 27,880,619.00 | 23,642,629.00 | 17,965,729.00 | 75.99 | 4,888.72 | 99.99 | 99.99 | 99.99 |

*All sequencing were performed using NovaSeq 6000

| Primerset ID | Enrichment method* | *T. pallidum* genome copies/µl in spiked samples (pre-enrichment) | $RNP_{Ct}$ in spiked samples (pre-enrichment) | *T. pallidum* genome copies/µl in spiked samples (post-enrichment) | $RNP_{Ct}$ in spiked samples (post-enrichment) |
|---|---|---|---|---|---|
| SWGA Pal 1 rep 1 | SWGA | 138.9 | 24.84 | 1,019,374 | 30.89 |
| SWGA Pal 1 rep 2 | SWGA | 111.8 | 25.1 | 826,351 | 30.94 |
| SWGA Pal 1 rep 3 | SWGA | 137.6 | 24.82 | 691,734 | 30.91 |
| SWGA Pal 2 rep 1 | SWGA | 138.9 | 24.84 | 1,013,850 | 27.29 |
| SWGA Pal 2 rep 2 | SWGA | 111.8 | 25.1 | 1,518,239 | 27.48 |
| SWGA Pal 2 rep 3 | SWGA | 137.6 | 24.82 | 1,421,354 | 27.79 |
| SWGA Pal 3 rep 1 | SWGA | 138.9 | 24.84 | 1,006 | 23.66 |
| SWGA Pal 3 rep 2 | SWGA | 111.8 | 25.1 | 841 | 23.32 |
| SWGA Pal 3 rep 3 | SWGA | 137.6 | 24.82 | 809 | 23.54 |
| SWGA Pal 4 rep 1 | SWGA | 138.9 | 24.84 | 6,884,365 | 28.35 |
| SWGA Pal 4 rep 2 | SWGA | 111.8 | 25.1 | 7,921,113 | 28.47 |
| SWGA Pal 4 rep 3 | SWGA | 137.6 | 24.82 | 5,731,638 | 28.5 |
| SWGA Pal 5 rep 1 | SWGA | 138.9 | 24.84 | 9,048,948 | 30.54 |
| SWGA Pal 5 rep 2 | SWGA | 111.8 | 25.1 | 10,391,119 | 30.33 |
| SWGA Pal 5 rep 3 | SWGA | 137.6 | 24.82 | 7,485,554 | 30.42 |
| SWGA Pal 6 rep 1 | SWGA | 138.9 | 24.84 | 57,838 | 25.39 |
| SWGA Pal 6 rep 2 | SWGA | 111.8 | 25.1 | 54,147 | 25.56 |
| SWGA Pal 6 rep 3 | SWGA | 137.6 | 24.82 | 40,579 | 25.74 |
| SWGA Pal 7 rep 1 | SWGA | 138.9 | 24.84 | 21,733 | 25.35 |
| SWGA Pal 7 rep 2 | SWGA | 111.8 | 25.1 | 29,324 | 25.13 |
| SWGA Pal 7 rep 3 | SWGA | 137.6 | 24.82 | 10,843 | 26.59 |
| SWGA Pal 8 rep 1 | SWGA | 138.9 | 24.84 | 181,057 | 25.38 |
| SWGA Pal 8 rep 2 | SWGA | 111.8 | 25.1 | 213,269 | 25.11 |
| SWGA Pal 8 rep 3 | SWGA | 137.6 | 24.82 | 181,281 | 25.1 |
| SWGA Pal 9 rep 1 | SWGA | 138.9 | 24.84 | 16,534,008 | 29.27 |
| SWGA Pal 9 rep 2 | SWGA | 111.8 | 25.1 | 14,923,347 | 29.25 |
| SWGA Pal 9 rep 3 | SWGA | 137.6 | 24.82 | 12,338,941 | 29.06 |
| SWGA Pal 10 rep 1 | SWGA | 138.9 | 24.84 | 13,545,202 | 28.63 |
| SWGA Pal 10 rep 2 | SWGA | 111.8 | 25.1 | 10,883,166 | 28.86 |
| SWGA Pal 10 rep 3 | SWGA | 137.6 | 24.82 | 10,244,179 | 28.9 |
| SWGA Pal 11 rep 1 | SWGA | 138.9 | 24.84 | 15,654,473 | 28.17 |
| SWGA Pal 11 rep 2 | SWGA | 111.8 | 25.1 | 13,267,096 | 28.23 |
| SWGA Pal 11 rep 3 | SWGA | 137.6 | 24.82 | 15,786,477 | 28.28 |
| SWGA Pal 12 rep 1 | SWGA | 138.9 | 24.84 | 9,212,052 | 27.54 |
| SWGA Pal 12 rep 2 | SWGA | 111.8 | 25.1 | 12,334,878 | 27.42 |
| SWGA Pal 12 rep 3 | SWGA | 137.6 | 24.82 | 9,056,400 | 27.32 |

*All sequencing were performed using NovaSeq 6000

| Sample/ Strain ID | Citation | SRA/NCBI_Accession/BioProject |
|---|---|---|
| AU15 | Arora_2016 | SRR3268722 |
| AU16 | Arora_2016 | SRR3268723 |
| BAL73 | Arora_2016 | SRR3268726 |
| CZ27 | Arora_2016 | SRR3268685 |
| NE15 | Arora_2016 | SRR3268705 |
| NE17 | Arora_2016 | SRR3268707 |
| NE19 | Arora_2016 | SRR3268709 |
| NE20 | Arora_2016 | SRR3268710 |
| SW6 | Arora_2016 | SRR3268732 |
| SW8 | Arora_2016 | SRR3268736 |
| DAL-1 | Čejková_2012 | NC_016844.1 |
| Nichols_v2 | Pětrošová_2013 | NC_021490.2 |
| Reference | Pětrošová_2013 | NC_021508.1 |
| SS14_v2 | Pětrošová_2013 | NC_021508.1 |
| PT_SIF0697 | Pinto_2016 | SRR3571774 |
| PT_SIF0751 | Pinto_2016 | SRR3584962 |
| PT_SIF0857 | Pinto_2016 | SRR3584965 |
| PT_SIF0877_3 | Pinto_2016 | SRR3571775 |
| PT_SIF0908 | Pinto_2016 | SRR3571776 |
| PT_SIF0954 | Pinto_2016 | SRR3571778 |
| PT_SIF1002 | Pinto_2016 | SRR3571783 |
| PT_SIF1020 | Pinto_2016 | SRR3571785 |
| PT_SIF1063 | Pinto_2016 | SRR3571786 |
| PT_SIF1135 | Pinto_2016 | SRR3584837 |
| PT_SIF1140 | Pinto_2016 | SRR3584838 |
| PT_SIF1142 | Pinto_2016 | SRR3584839 |
| PT_SIF1156 | Pinto_2016 | SRR3584840 |
| PT_SIF1167 | Pinto_2016 | SRR3584841 |
| PT_SIF1183 | Pinto_2016 | SRR3584842 |
| PT_SIF1196 | Pinto_2016 | SRR3584843 |
| PT_SIF1200 | Pinto_2016 | SRR3584844 |
| PT_SIF1242 | Pinto_2016 | SRR3584845 |
| PT_SIF1252 | Pinto_2016 | SRR3584871 |
| PT_SIF1261 | Pinto_2016 | SRR3584875 |
| PT_SIF1278 | Pinto_2016 | SRR3584879 |
| PT_SIF1280 | Pinto_2016 | SRR3584882 |
| PT_SIF1299 | Pinto_2016 | SRR3584884 |
| B3 | Sun_2016 | SRR2996728 |
| C3 | Sun_2016 | SRR2996729 |
| K3 | Sun_2016 | SRR2996730 |
| Q3 | Sun_2016 | SRR2996732 |
| SHC-0 | Sun_2016 | SRR2996724 |
| SHD-R | Sun_2016 | SRR2996725 |
| SHE-V | Sun_2016 | SRR2996726 |

| | | |
|---|---|---|
| SHG-I2 | Sun_2016 | SRR2996727 |
| Chicago_tube2 | Beale_2019 | ERS2676913 |
| Mexico_A | Beale_2019 | NC_018722.1 |
| Nichols_Houston_E | Beale_2019 | ERS2676914 |
| Nichols_Houston_J | Beale_2019 | ERS2676912 |
| Nichols_Houston_O | Beale_2019 | ERS2676915 |
| Seattle_81-4 | Beale_2019 | ERS2676916 |
| UW074B | Beale_2019 | ERS1884608 |
| UW093B | Beale_2019 | ERS1884633 |
| UW099B | Beale_2019 | ERS1884579 |
| UW102B | Beale_2019 | ERS1884607 |
| UW104B | Beale_2019 | ERS1884521 |
| UW116B | Beale_2019 | ERS1884567 |
| UW125B | Beale_2019 | ERS1884612 |
| UW134B | Beale_2019 | ERS1884507 |
| UW138B | Beale_2019 | ERS1884616 |
| UW140B | Beale_2019 | ERS1884597 |
| UW147B | Beale_2019 | ERS1884615 |
| UW148B | Beale_2019 | ERS1884619 |
| UW149B | Beale_2019 | ERS1884519 |
| UW155B | Beale_2019 | ERS1884511 |
| UW181B | Beale_2019 | ERS1884618 |
| UW187B | Beale_2019 | ERS1884605 |
| UW195B | Beale_2019 | ERS1884514 |
| UW211B | Beale_2019 | ERS1884629 |
| UW213B | Beale_2019 | ERS1884621 |
| UW215B | Beale_2019 | ERS1884631 |
| UW228B | Beale_2019 | ERS1884606 |
| UW231B | Beale_2019 | ERS1884627 |
| UW244B | Beale_2019 | ERS1884505 |
| UW254B | Beale_2019 | ERS1884603 |
| UW254C | Beale_2019 | ERS1884626 |
| UW259B | Beale_2019 | ERS1884506 |
| UW262B | Beale_2019 | ERS1884572 |
| UW264B | Beale_2019 | ERS1884518 |
| UW280B | Beale_2019 | ERS1884512 |
| UW281B | Beale_2019 | ERS1884628 |
| UW291B | Beale_2019 | ERS1884580 |
| UW298B | Beale_2019 | ERS1884552 |
| UW303B | Beale_2019 | ERS1884520 |
| UW304B | Beale_2019 | ERS1884515 |
| UW327B | Beale_2019 | ERS1884516 |
| UW330B | Beale_2019 | ERS1884613 |
| UW337B | Beale_2019 | ERS1884573 |
| UW344B | Beale_2019 | ERS1884602 |
| UW368B | Beale_2019 | ERS1884609 |
| UW370B | Beale_2019 | ERS1884557 |

| | | |
|---|---|---|
| UW376B | Beale_2019 | ERS1884585 |
| UW383B | Beale_2019 | ERS1884625 |
| UW391B | Beale_2019 | ERS1884576 |
| UW391C | Beale_2019 | ERS1884591 |
| UW411B | Beale_2019 | ERS1884604 |
| UW473B | Beale_2019 | ERS1884596 |
| UW492B | Beale_2019 | ERS1884636 |
| UW526B | Beale_2019 | ERS1884508 |
| UW823B | Beale_2019 | ERS1884588 |
| UW824B | Beale_2019 | ERS1884558 |
| UW852B | Beale_2019 | ERS1884620 |
| NL10 | Beale_2019 | ERS1724924 |
| NL11 | Beale_2019 | ERS1724925 |
| NL13 | Beale_2019 | ERS1724927 |
| NL14 | Beale_2019 | ERS1724928 |
| NL16 | Beale_2019 | ERS1724930 |
| NL17 | Beale_2019 | ERS1724931 |
| NL19 | Beale_2019 | ERS1724933 |
| Amoy | Tong 2017 | NZ_CP015162.1 |
| CDC-A | unpublished | NZ_CP010559.1 |
| Chicago_Population | unpublished | NZ_CP010558.1 |
| Seattle_Nichols | unpublished | NZ_CP010422.1 |
| CW30 | Grillova_2019 | GCA_005311165.1 |
| CW56 | Grillova_2019 | GCA_005311145.1 |
| CW59 | Grillova_2019 | GCA_005311445.1 |
| CW65 | Grillova_2019 | GCA_005311125.1 |
| CW82 | Grillova_2019 | GCA_005311105.1 |
| CW83 | Grillova_2019 | GCA_005311085.1 |
| CW84 | Grillova_2019 | GCA_005311205.1 |
| CW85 | Grillova_2019 | GCA_005311185.1 |
| CW86 | Grillova_2019 | GCA_005311065.1 |
| CW87 | Grillova_2019 | GCA_005341485.1 |
| CW88 | Grillova_2019 | GCA_005311045.1 |
| Grady | Grillova_2019 | GCA_005408405.1 |
| Philadelphia-1 | Grillova_2019 | GCA_005408385.1 |
| Nichols-CDC | This study | PRJNA744275 |
| CDC-SF003 | This study | PRJNA744275 |
| EUHM-004 | This study | PRJNA744275 |
| EUHM-012 | This study | PRJNA744275 |
| EUHM-013 | This study | PRJNA744275 |
| EUHM-014 | This study | PRJNA744275 |
| STLC-001 | This study | PRJNA744275 |
